# Comparative transcriptomics of immune response to viral and bacterial stimuli in three acanthopterygian bony fish

**DOI:** 10.64898/2026.01.30.702270

**Authors:** Raquel Rodríguez-Vázquez, Manu Kumar Gundappa, Oscar Aramburu, Jelena Radojicic, Costas S. Tsigenopoulos, Serena Ferraresso, Raffaella Franch, Luca Bargelloni, Paulino Martínez, Diego Robledo, Hendrik-Jan Megens

## Abstract

Diseases triggered by bacterial and viral infections have caused huge economic losses for three of the most important European aquaculture species: turbot (*Scophthalmus maximus*), gilthead seabream (*Sparus aurata*) and European seabass (*Dicentrarchus labrax*). Understanding how they respond to pathogens is relevant for advancing aquaculture disease management and comprehending evolution of immune response within teleosts. Since mechanisms conserved across species are assumed to perform important roles, comparative analysis provides a powerful approach to pinpoint key elements of the immune defence. Here, we report the first comparative immune-transcriptomic analysis of these three species using bacterial and viral mimics after 20-24 hours post-stimulation with inactivated *Vibrio anguillarum* and Poly I:C in the head kidney of live fish (*in vivo*), and in primary leukocyte cultures (*in vitro*). The transcriptomic response, based on RNA-seq data, revealed a total of 503 differentially expressed orthologous genes in response to *in vitro*-Poly I:C, 1,472 to *in vitro*-*Vibrio*, 920 to *in vivo*-Poly I:C, and 832 to *in vivo*-*Vibrio*. Interestingly, consistent expression patterns were identified in seven genes across all species in both cell culture and live organisms in response to both pathogen stimuli. Functional enrichment analysis revealed associations with immunity, DNA replication and repair, and cytokine pathways, with the Toll-Like Receptor (TLR) pathway common to both conditions and stimuli. Our study suggests conservation of orthologous gene expression during infection across the three species for genes involved in chemokine pathways, interferon signalling, antigen processing and presentation, cell signalling regulators, and MAPK cascades. This study provides insights into key immune defence mechanisms in acanthopterygian bony fish.

## 1. INTRODUCTION

Aquaculture is of crucial importance in addressing the growing global demand for marine products [1], particularly in Europe, where species such as turbot (*Scophthalmus maximus*), gilthead seabream (*Sparus aurata*) and European seabass (*Dicentrarchus labrax*) are highly appreciated for their rapid growth and production efficiency [2–5]. However, with the growth of the aquaculture industry, bacterial and viral diseases have become increasingly severe and result in substantial economic losses [6].

During infection, a complex interaction takes place between hosts and pathogens, whereby the host immune system activates genes within immune pathways aimed at neutralizing the infection. Fish depend significantly on their innate immunity, which serves as their primary defence mechanism. Physical barriers such as skin, scales and mucosal surfaces provide physical immunity. Additionally, these mucosal surfaces also confer mucosal immunity through secretory immunoglobulins, which work alongside antimicrobial peptides (AMPs) capable of disrupting microbial membranes [7]. Furthermore, fish possess pattern recognition receptors (PRRs) that recognize pathogen-associated molecular patterns (PAMPs) such as key structural components of bacterial cell walls (e.g., lipopolysaccharides (LPS) and peptidoglycans) and viral RNA. These receptors initiate signalling pathways that activate pro-inflammatory responses in fish, which include cytokines that act as messengers to start inflammation, recruit immune cells, and activate antimicrobial defence mechanisms. The head kidney is essential for haematopoiesis and the production of cytokines, while the spleen contributes to the filtration of pathogens and the coordination of inflammatory responses. Concerning adaptive immunity, while it is not as developed as in mammals, fish possess T and B cells that generate specific antibodies (mainly IgM and IgT) to neutralize pathogens [8].

The great diversity of aquaculture fish species and infectious pathogens poses a challenge for unravelling immune response pathways. However, it can be assumed that immune responses include conserved pathways, and that different fish species will present similarities in their response to, for example, bacterial or RNA-viral infections. Therefore, comparative immunology represents an efficient tool to comprehend the main mechanisms underpinning immune responses in a group as diverse as fish [9,10]. Comparative research on the immune response in fish to various pathogens has made significant advances in recent years, revealing complex interactions between aquatic species and infectious agents. Notably, comparative analyses are often challenged by the fact that most pathogens are either species-specific or exhibit species-specific serotypes, making direct comparisons difficult [11]. To overcome this issue, recent studies have employed both PAMPs, such as Poly I:C (Polyinosinic:polycytidylic acid) or LPS, and generalist pathogens, such as viral haemorrhagic septicaemia, *V. anguillarum* or *Streptococcus parauberis* [12–14]. The use of PAMPs provides an effective method for comparing immune responses across species, without the effect of pathogen specificity, thereby enabling more robust interspecies comparisons [15]. In aquaculture species, a considerable number of transcriptomic studies have been conducted with the objective of elucidating the immune response of fish to inactivated or alive *V. anguillarum* [16–18] or to dsRNA viruses using Poly I:C [19–23], representing common pathogens in farmed settings. In addition to comparisons between species, the difference between *in vivo* and *in vitro* conditions is also of interest due to the reduced complexity of cell culture models and widespread use to study infectious diseases. Saravia et al. (2022) [24] studied the transcriptomic response of *Harpagifer antarcticus* to LPS and Poly I:C both *in vitro* and *in vivo*, while Aramburu et al. (2025) [23] assessed not only the turbot transcriptome following exposure to Poly I:C and *V. anguillarum* both *in vivo* and *in vitro*, but also epigenetic regulation using ATAC-seq and ChIP-seq. However, a review of the current scientific literature reveals no studies comparing the immune responses of different fish species, to Poly I:C and bacteria, while also considering both *in vitro* and *in vivo* conditions.

Unravelling the adjustments of gene expression in the host is fundamental for the comprehension of the infection process at the molecular level to develop strategies for disease control [25]. In recent years, transcriptome analysis using high-throughput RNA sequencing (RNA-seq) has emerged as a powerful tool for elucidating the immune response mechanisms of fish, understanding the molecular basis of pathogen resistance, and examining the comparative immune response across species [16,26]. A comparative functional annotation of immune responses to two categories of disease pathogens (such as bacterial and viral) entails the identification and description of the roles played by genes throughout the entire genome. As a part of the European AQUA-FAANG project, we conducted an RNA-seq analysis of head kidney samples from turbot, seabream and seabass after exposure to heat-killed *V. anguillarum* and Poly I:C treatment, using both the full organ (*in vivo*) and primary leukocyte cultures (*in vitro*). The objectives of this study were first to evaluate anti-viral and anti-bacterial responses by direct stimulation of primary leukocyte cultures in comparison with the head kidney of intraperitoneal injected fish, and secondly, to identify conserved immune response pathways by comparing expression in orthologous genes across the three species. To our knowledge, this is the first comparative immune response in these species, and the findings would provide new tools for improving disease resistance in aquaculture.

## 2. MATERIAL AND METHODS

### 2.1 Animals

Thirty specimens of *S. maximus,* twenty-six of *S. aurata* and twenty-four of *D. labrax* were used for the study (Table S1). Sampling followed the AQUA-FAANG protocols https://data.faang.org/api/fire_api/experiments/INRA_SOP_invivo.invitro.challenges_20200131.pdf and details are provided in Supplementary Methods, Tables S1 and S2.

### 2.2 In vivo immunostimulation

Eighteen fish were intraperitoneally injected per species with Poly I:C (viral mimic, 6 replicates), heat-killed *V. anguillarum* (bacterial stimulus, 6 replicates), or PBS (control, 6 replicates). Before inoculation, Poly I:C (Sigma P1530; 5 mg/ml in PBS) was heated at 55 °C for 15 min, cooled to room temperature (20 min) and administered at five µg/g of body weight. *V. anguillarum* (strain P0382; INRA, France) was cultured in tryptic soy broth to an OD600 of 1.5. The pellet from 100 ml culture was washed four times with isotonic NaCl (9 g/L), resuspended in 1 ml, heat-inactivated at 100°C for 30 seconds, cooled, and stored at −80°C. Each specimen received 138 µl of 1:10 PBS-diluted bacterial extract. Controls were administered 100 µl of PBS. Following a 20-24 h period, fish were anesthetized by bath (MS-222; 100 mg/L) and euthanized with an anaesthetic overdose (MS-222; 150 mg/L), and head kidney samples were extracted, washed with PBS, cut (>20 mg each fragment), and either flash frozen in liquid nitrogen or preserved in RNAlater (Thermofisher Scientific) for downstream RNAextraction and sequencing protocols. Thereafter, all samples were stored at −80°C (Fig. 1A, Tables S1 and S2).

**Fig. 1.**
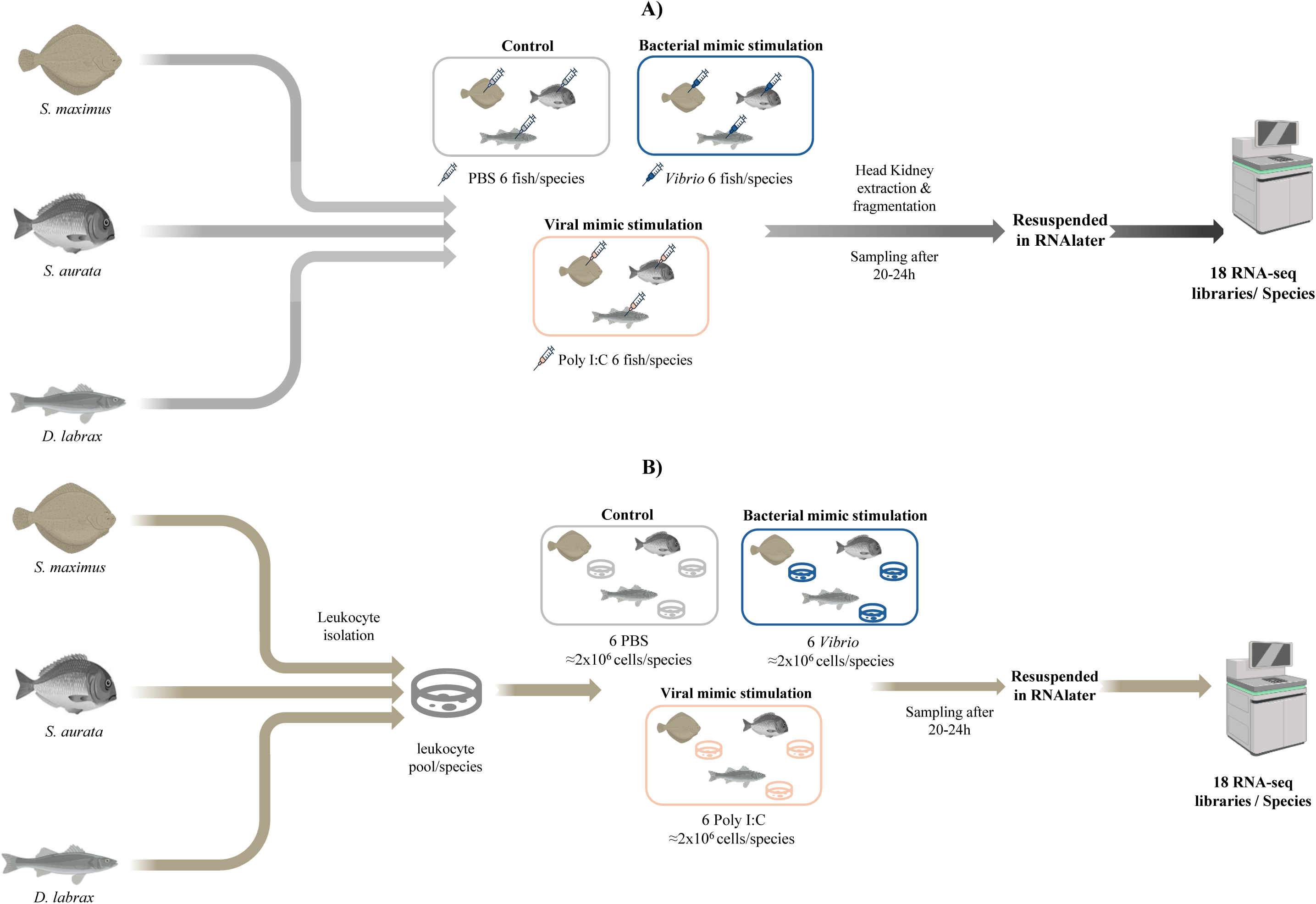
Experimental workflow for *S. maximus*, *S. aurata* and *D. labrax* stimulations: A) *In vivo* stimulation with PBS, Poly I:C, and *Vibrio*, followed by subsequent sampling of head kidney; B) *In vitro* stimulation with head kidney leukocyte cultures using the same agents.

### 2.3 In vitro immunostimulation

Leukocytes were isolated from turbot, seabream, and seabass fish. The entire head kidney was aseptically isolated and placed in Petri dishes containing 40 ml of cell isolation media (500 ml of Leibovitz L-15 medium (L-15), 10 ml FBS (2%), and 0.02% EDTA. Subsequently, samples were cut and filtered through a 100 µm nylon mesh at a constant flow of cell isolation media. Leukocytes were separated by centrifugation (400 x g, 10 min, 4°C) of a 40-ml cell suspension that was layered in a 50 ml tube containing 51% Percoll. The interface layer was collected, centrifuged (400 x g, 10 min, 4°C), and washed three times with L-15 medium with 0.1% FBS. After cell counting and viability assessment (trypan blue exclusion test), samples were pooled and divided into aliquots of 2 x 10⁶ cells to reach six replicates per treatment. Each of six technical replicates was stimulated with 20 µl of Poly I:C solution, 20 µl of inactivated *V. anguillarum,* and 20 µl PBS (controls). Leukocyte cultures were incubated for 20-24 h at 16 °C. Cells were collected in 2 ml Eppendorf tubes, pelleted (500 × g for 5 minutes at room temperature), resuspended in RNAlater or flash frozen in liquid nitrogen and stored at −80 °C for RNA-seq (Fig. 1B, Tables S1 and S2).

### 2.4 RNA isolation and sequencing

RNA extraction was performed according to the FAANG protocols for both *in vivo* and *in vitro* stimulations (data.faang.org; see Table S2). Briefly, RNA was isolated and purified using the miRNeasy Kit (QIAGEN) with specific protocol adjustments for head kidney (> 20 mg) and isolated leukocytes (2 × 10^6^ cells). RNA quality and quantity were assessed using a Bioanalyzer (Bonsai Technologies, Madrid, Spain) and a NanoDrop® ND-1000 spectrophotometer (NanoDrop® Technologies Inc., Wilmington, DE, USA). Libraries were prepared at Novogene (UK) using NEBNext Ultra Directional RNA Library Prep Kits (Illumina). Sequencing was performed on an Illumina NovaSeq S4 platform, producing 150 bp paired end reads.

### 2.5 Differential expression analysis

Read quality was assessed using FastQC (v0.12.1) [27] and raw reads were trimmed using FastP v0.22.0, with Phred quality <15 and reads with length <30 bp [28]. Clean reads were pseudo-aligned to the corresponding reference transcriptome using Kallisto (v0.46.1) [29] and gene expression was quantified using 100 bootstraps. The reference transcriptomes used were ASM1334776v1 (GCA_013347765.1) for *S. maximus*, fSpaAur1.1(GCA_900880695.1) for *S. aurata* and dlabrax2021 (GCA_905237075.1) for *D. labrax*. Count data were filtered to remove genes with fewer than 5 reads in all samples and represented in only one sample across all conditions. Raw counts were analysed in R v4.3.3 using DESeq2 [30] for differential expression. Variance stabilizing transformation was applied to normalize counts prior to quality assessment and identification of potential outliers through Principal Component Analysis (PCA) plots. Genes with a false discovery rate (FDR) adjusted p-value less than 0.05 were identified as differentially expressed genes (DEGs). Analysis in *S. maximus*, *S. aurata* and *D. labrax* data were all performed using the same pipeline.

### 2.6 Enrichment and gene ontology analysis

Functional enrichment of the lists of DEGs was performed using g:Profiler (version e111_eg58_p18_f463989d) [31] using the Ensembl reference genomes mentioned in Section 2.5 (last accession in October, 2024). Enriched biological process Gene Ontology (GO) terms were estimated with the Benjamini-Hochberg FDR adjusted *P*-value < 0.05 in Reduce + Visualize Gene Ontology software (ReviGO v1.8.1) [32], applying SimRel semantic similarity (threshold: 0.4) to reduce redundancy. Immune-related GO terms were identified using the online tools QuickGO (https://www.ebi.ac.uk/QuickGO/annotations) [33] and AmiGO2 (https://amigo.geneontology.org/amigo) [34] (Last accession on October, 2024). Immune-related DEGs were screened and orthologues among species were identified using the Ensembl Biomart tool via the BiomaRt package (v2.58.2 in R) [35,36]. The resulting orthology mapping table was used to map immune-related DEGs across these species, facilitating comparative analysis of immune gene expression patterns. Venn Diagrams [37] and heatmaps (pheatmap v1.0.12, scale: “row” in R) [38] were used to visualize DEG distributions and expression patterns.

### 2.7 Identification and functional analysis of conserved orthologous genes

To infer the homology of response in the three species using DEGs, five lists of potentially conserved genes were compiled. Each list included DEGs in at least one species with consistent regulatory trends in the others, defined as genes showing the same upregulation or downregulation across all three species. List 1 contained genes consistently responding to both Poly I:C and *Vibrio* under *in vitro* and *in vivo* conditions. Lists 2 and 3 included genes with conserved responses *in vitro* or *in vivo* conditions and lists 4 and 5 contained genes with conserved response either to Poly I:C, or *Vibrio* challenges. The latter four lists were further annotated by Kyoto Encyclopedia of Genes and Genomes (KEGG) pathway analysis [39] using the Database for Annotation, Visualization and Integrated Discovery (DAVID) tool [40]. For species comparison, 1:1 orthologs to *S. maximus* were used to identify the biological pathways that were significantly enriched. The KEGG pathway enrichment analysis was performed with FDR < 0.05. KEGG Mapper was used to provide a graphic representation of interactions between genes. To further investigate the relationship between genes with the same expression profile, protein interaction networks were constructed using STRING v12.0 for the lists [41] based on the *S. maximus* proteome annotation.

## 3. RESULTS

### 3.1. Sample metadata

A total of 108 RNA-seq datasets were used in this study: 36 for turbot, 36 for seabream and 36 for seabass. These samples represented 6 experimental groups: control (mock-challenged with PBS), challenged with *V. anguillarum* and challenged with Poly I:C, both *in vivo* and *in vitro*. Full sample and metadata information is provided in Tables S1 and S2.

### 3.2. RNA-sequencing

For turbot, the average number of RNA-seq raw reads per sample was 69,606,581, with an average mapping of 98.9% to the turbot transcriptome (Table S3A); for seabream, the average number of raw reads was 66,673,811 and 99.1% mapping to its transcriptome (Table S3B); and for seabass 67,880,066 raw reads on average and 99.2% mapping to its transcriptome (Table S3C).

Clear PCA distinctions are evident between control and stimulated samples across species and conditions (Fig. S1). In *in vitro* conditions, Poly I:C stimulation resulted in the strongest separation in *S. aurata* and the lowest in *D. labrax*, while *Vibrio* stimulation showed consistent differentiation across species. In *in vivo*, both Poly I:C and *Vibrio* stimulations demonstrated non-overlapping groups, with clear separations along PC1 and PC2 (see details in Supplementary Results).

A differential expression analysis was conducted to compare each stimulated condition with its respective control for both *in vitro* and *in vivo* conditions. Overall, 55.03% of the genes were upregulated in the treatments compared to the controls, and 49.96% were downregulated on average (Fig S2A). Moreover, the *in vitro Vibrio* stimulation revealed a significant and somewhat similar number of DEGs in all three species compared to the other experimental conditions with 8,783, 6,238, and 7,296 DEGs in turbot, seabream and seabass, respectively (Fig. S2B, Table S4). In the other three conditions, the response was quite species-specific, with seabream showing the weakest response in the two *in vivo* conditions (Poly I:C (856 DEGs) and *Vibrio* (430 DEGs)), while the strongest one was in the *in vitro* Poly I:C (4,335 DEGs); turbot showed the greater number of DEGs in *in vivo* Poly I:C (6,101 DEGs), compared to *in vitro* Poly I:C (1,383 DEGs), and *in vivo Vibrio* (2,557 DEGs); finally, seabass showed a strong response *in vivo* both for *Vibrio* (7,296 DEGs) and Poly I:C (6,586 DEGs), while it showed a very weak for *in vitro* Poly I:C (504 DEGs). In general, turbot showed a slightly higher *in vitro* than *in vivo* response (10,166 vs 8,658 DEGs), seabream showed the same trend but with a much larger difference (10,573 vs 1,286 DEGs), and seabass showed the opposite trend, with a higher number of DEGs *in vivo* than *in vitro* response (13,881 vs 7,800 DEGs). Additionally, the UpSet plot analysis revealed that shared DEGs across conditions within each species displayed species-specific patterns (Fig. S3). In *S. aurata*, the highest number of shared DEGs (2,619) was observed under *in vitro* conditions (Poly I:C and *Vibrio* exposures). For *S. maximus*, 1,862 shared DEGs were found in the comparison between the *in vitro* (*Vibrio*) and *in vivo* (Poly I:C) experiments. A high number of shared DEGs was also observed between *in vivo* (Poly I:C and *Vibrio*) and *in vitro* (*Vibrio*) challenges for *D. labrax* (2,786).

### 3.3 Functional enrichment among DEGs

Biological Processes (BP) Gene Ontology (GO) analysis of DEGs in the three teleost species revealed conserved immune-related pathways across viral (Poly I:C) and bacterial (*Vibrio*) challenges, with notable differences in species-specific responses. REVIGO-clustered GO terms highlighted shared immune mechanisms, such as immune system process (GO:0002376), immune response (GO:0006955), response to external stimuli/pathogens (e.g., GO:0006952; GO:0009615), and interspecies interaction (GO:00044419), among others. These results provide a framework for comparative transcriptomic studies of immune evolution in teleost fish (see details in Supplementary Results, Tables S5, S6 and Figure S4).

### 3.4 Turbot, Seabream and Seabass head kidney comparative transcriptome

A comparative analysis of immune response-associated DEGs across the three species revealed a small but consistent set of shared genes in each challenge, while most responses remained species-specific across both *in vitro* and *in vivo* with Poly I:C and *Vibrio* stimulations. Notably, *in vitro* stimulation with Poly I:C revealed upregulation in 31 shared genes across the three species. Overall, turbot exhibited a more divergent transcriptional profile, with stronger induction of interferon and immune signalling genes, whereas seabream and seabass showed higher expression of genes involved in antigen processing and cell regulation (see detailed in Supplementary Results; Table S7, Fig. 2).

**Fig 2.**
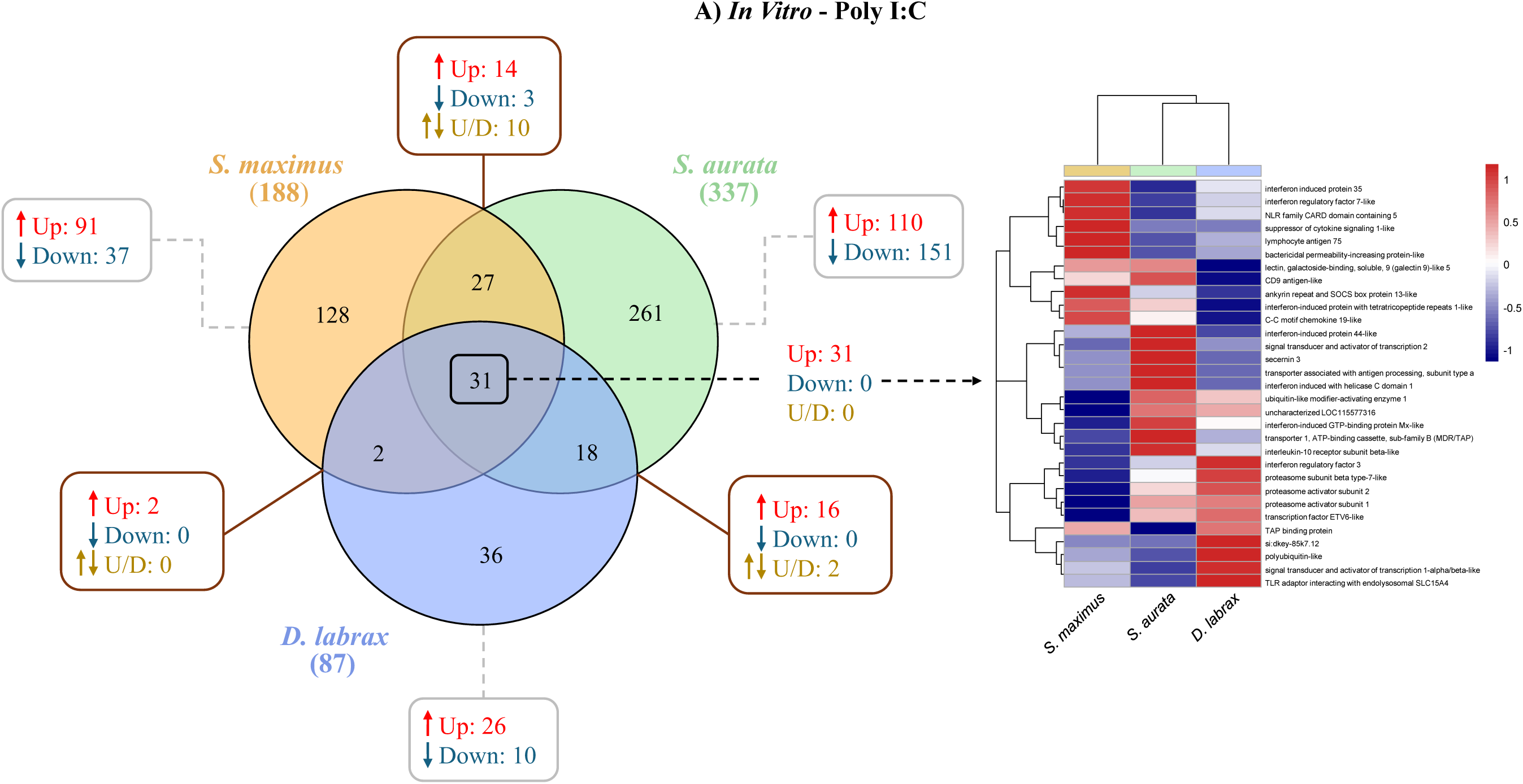
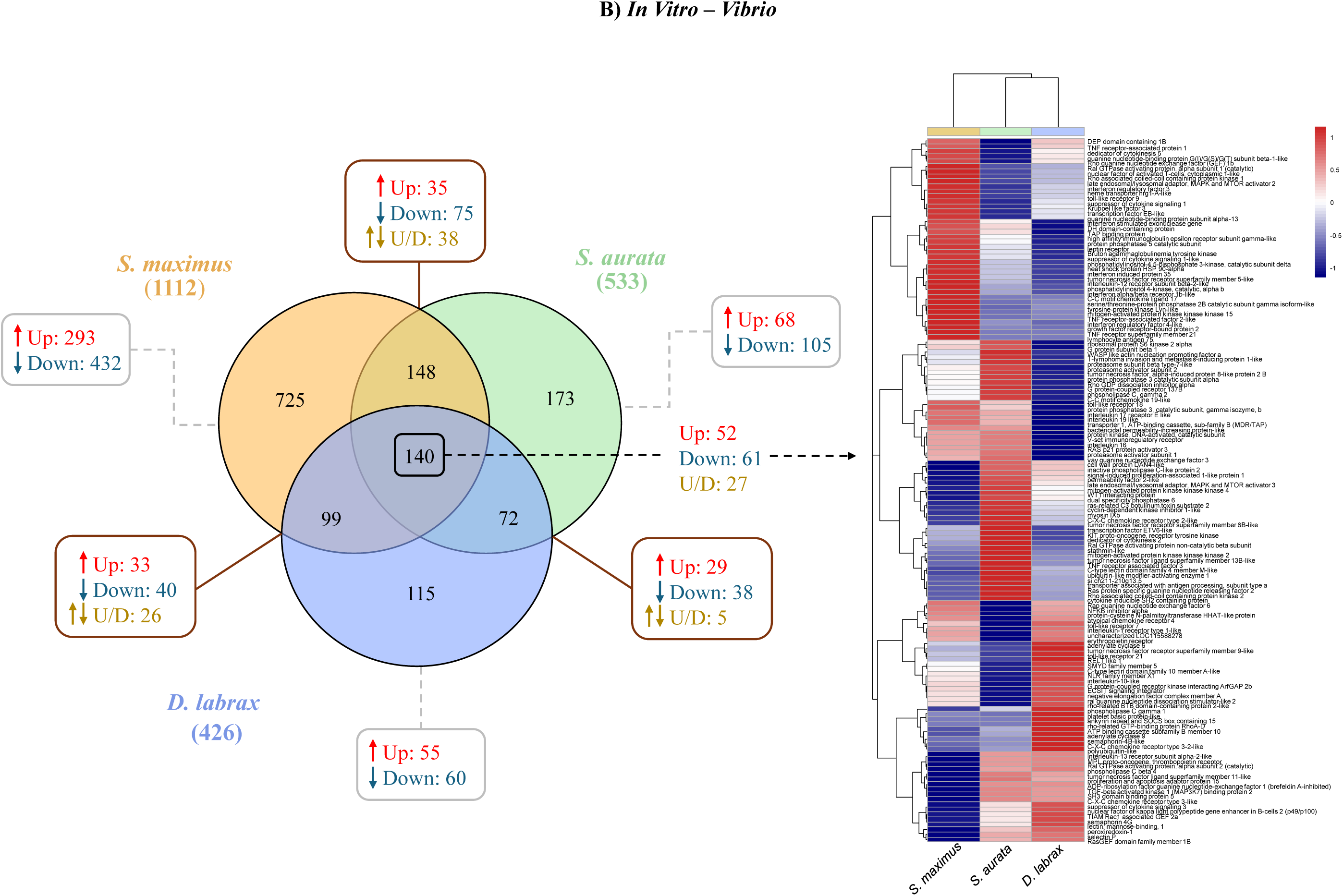
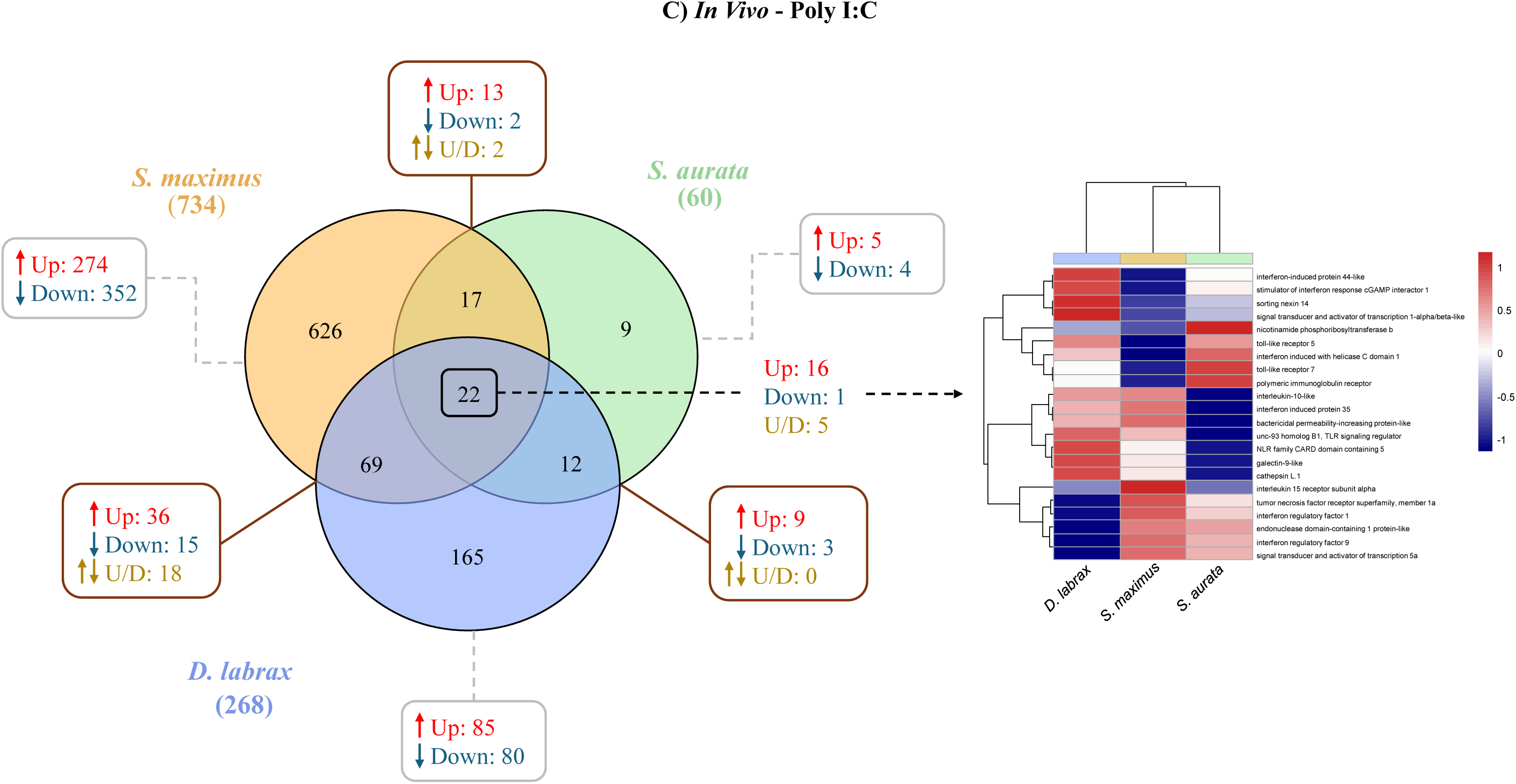
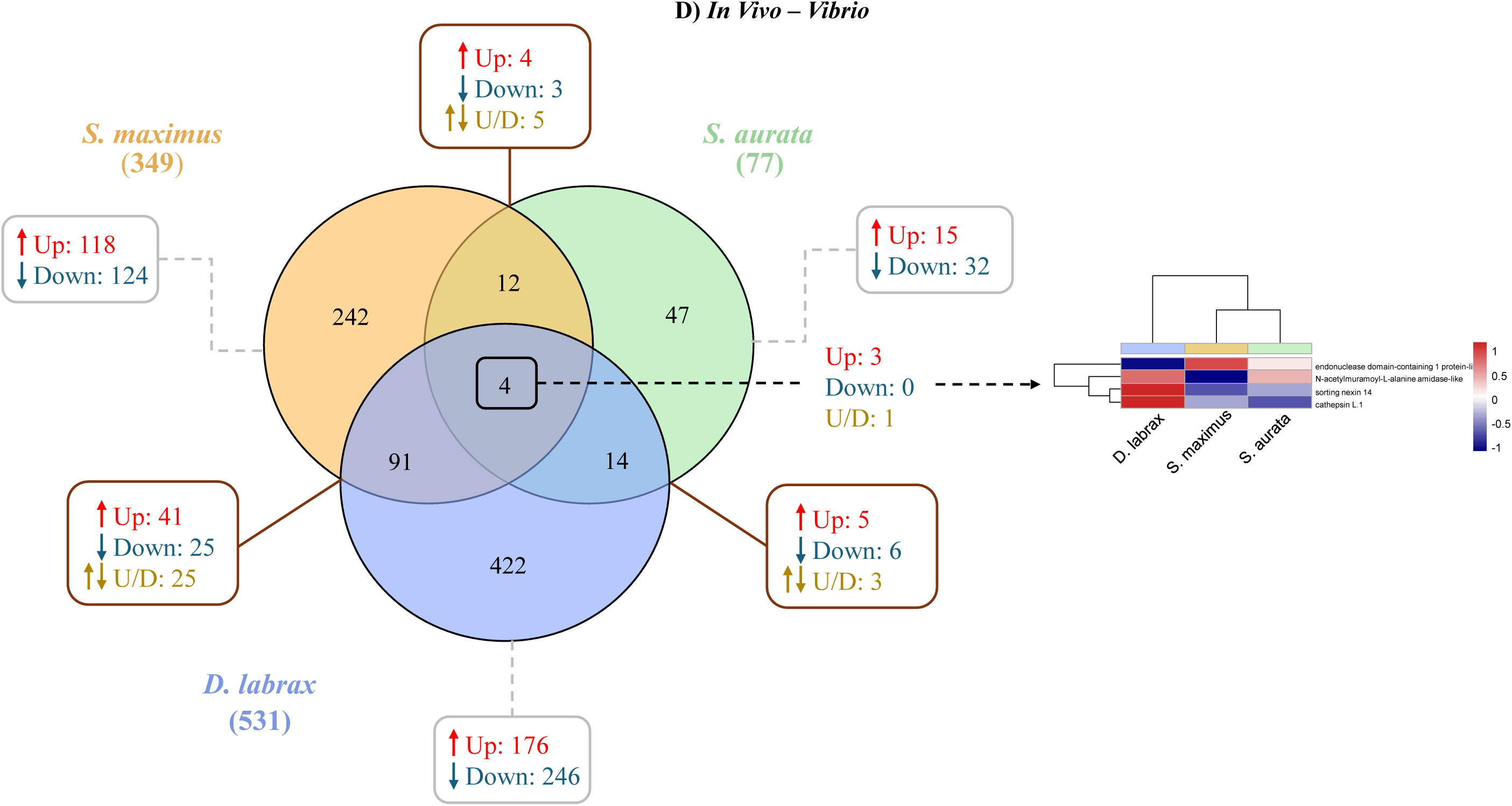
Venn’s diagrams showing the unique and common differentially expressed genes (DEGs) among *S. maximus*, *S. aurata* and *D. labrax* and heatmaps of expression-normalized DEGs across species. A) *In vitro* response to Poly I:C stimulation, B) *In vitro* response to *Vibrio* stimulation, C) *In vivo* response to Poly I:C stimulation, and D) *In vivo* response to *Vibrio* stimulation. Venn’s diagrams illustrate genes that are up-(red), down-(blue), or up/down-regulated (yellow) between different species. Color scale in heatmaps: −1 (blue, below mean) to 1 (red, above mean), representing deviations from the average expression.

### 3.5 Conserved immune responses across fish species

We identified conserved genes activated by Poly I:C across *in vitro* and *in vivo* models (Table S8) and *Vibrio-*Poly I:C challenges in cell cultures of the three species (Table S9). Seven DEGs were consistently upregulated (*bpifcl, ifi35, ifi44, ifih1, lgals9l, nlrc5, stat1a*) in both cell cultures and live fish following exposure to Poly I:C, but not in response to *Vibrio* (see detailed Suppl. Results, Table S8); 15 DEGs were detected across the three species when challenged *in vitro* with both Poly I:C and *Vibrio,* with 13 exhibiting the same directional expression change (*ccl19, ifi35, irf3, isg15, psme1, psme2, psmb13a, socs1b, tapbp2, etv7, tap1, tap2a, uba7*). Notably, *bpifcl* was upregulated against Poly I:C but downregulated against *Vibrio*, while *ly75* displayed species-specific regulation. No overlapping DEGs were detected *in vivo* for *Vibrio* and Poly I:C (see details in Suppl. Results, Table S9).

Moreover, DEGs in at least one species which showed the same direction of change in the other two species were identified to look for consistent patterns of gene regulation (Table S10A). A total of 7 genes were detected in all species and challenges, four upregulated: endoplasmic reticulum protein 44 (*erp44*), chemokine (C-C motif) ligand 19 (*ccl19*), proteasome 20S subunit beta 9a (*psmb9a*), immunoglobin binding protein 1 (*igbp1*), while three were downregulated: mitogen-activated protein kinase kinase 6 (*map2k6*), IL2 inducible T cell kinase (*itk*) and arrestin beta 2 (*arrb2*). An interaction network was explored between those genes using the STRING program, which showed that all genes were interconnected except for *igbp1*. It was also observed that *itk* was related to all nodes, showing a weaker interaction with the *errp44* gene, but a strong interaction with the remaining genes (Fig. 3).

**Fig 3.**
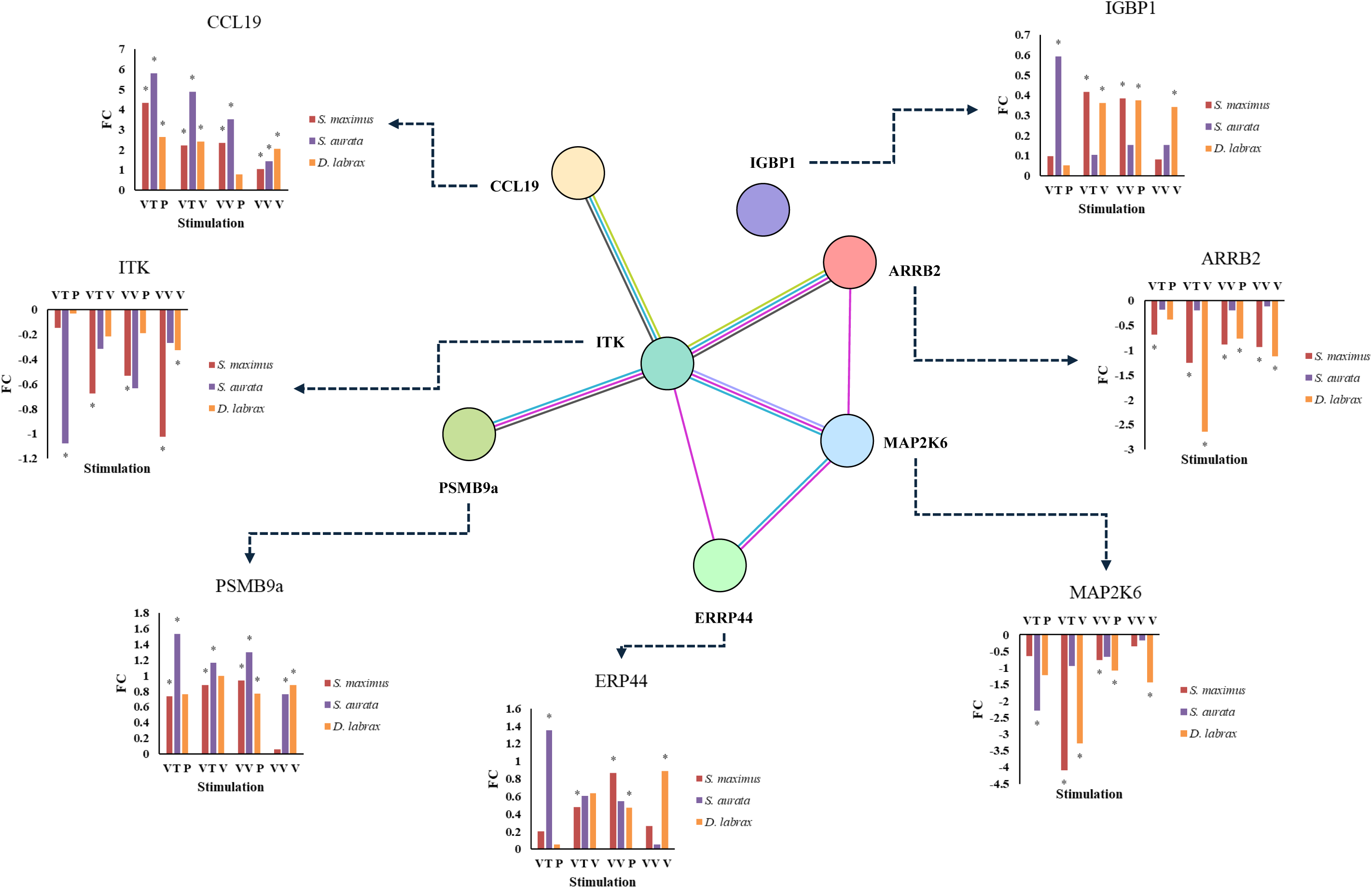
Protein-protein interaction (PPI) network based on the STRING database and showing fold change profiles. The interaction pattern and *fold change* (*FC*) were inspected in differentially expressed genes identified in at least one of the three species while maintaining consistent regulation direction across all species under the four stimulation scenarios. The network was performed with a confidence score of 0.150. Nodes representing genes and lines between them are the edges, indicating types of evidence for interactions. Blue and pink edges are known interactions (from curated databases and experimentally determined, respectively); grey, green and purple edges are interactions derived from text mining, co-expression and protein homology respectively. Gene expression changes across species and challenges are described in bar charts. X-axis represents experimental conditions: VT P (*In Vitro*-Poly I:C), VT V (*In Vitro-Vibrio*), VV P (*In Vivo*-Poly I:C), and VV V (*In Vivo-Vibrio*). Y-axis: *FC* between control and stimulated conditions. Asterisks (*) indicate significant differences (p < 0.05) between control and stimulated conditions for each species in individual experiments (Table S10A).

In addition, genes that shared similar regulatory patterns were also analysed separately in the *in vitro* and *in vivo* conditions and under Poly I:C and *Vibrio* stimulations. The list of identified genes is presented in Table S10. For *in vitro*, a total of 140 genes showed the same pattern for Poly I:C and *Vibrio* (Table S10B), while i*n vivo* a total of 59 genes were detected (Table S10C). Meanwhile for Poly I:C a total of 54 genes were revealed (Table S10D) and under *Vibrio* stimulation a total of 90 genes showed the same pattern (Table S10E).

Enrichment pathway analyses for each of these lists were conducted using the Kyoto Encyclopaedia of Genes and Genomes (KEGG) database (see detailed in Suppl. Results, Table S11, Fig. S5). Analyses demonstrated that the toll-like receptor (TLR) signalling pathway is a common pathway of all experimental conditions (Fig. S5). A detailed analysis of the TLR signalling pathway reveals a complex network of interactions involving several interconnected signalling pathways for *in vitro* (Fig. 4A, Table S12A) and *in vivo* (Fig. 4C, Table S12B), as well as under Poly I:C (Fig. 5A; Table S12C) and *Vibrio* (Fig 5C, Table S12D) stimulations. Further details are provided in Supplementary Results.

**Fig. 4.**
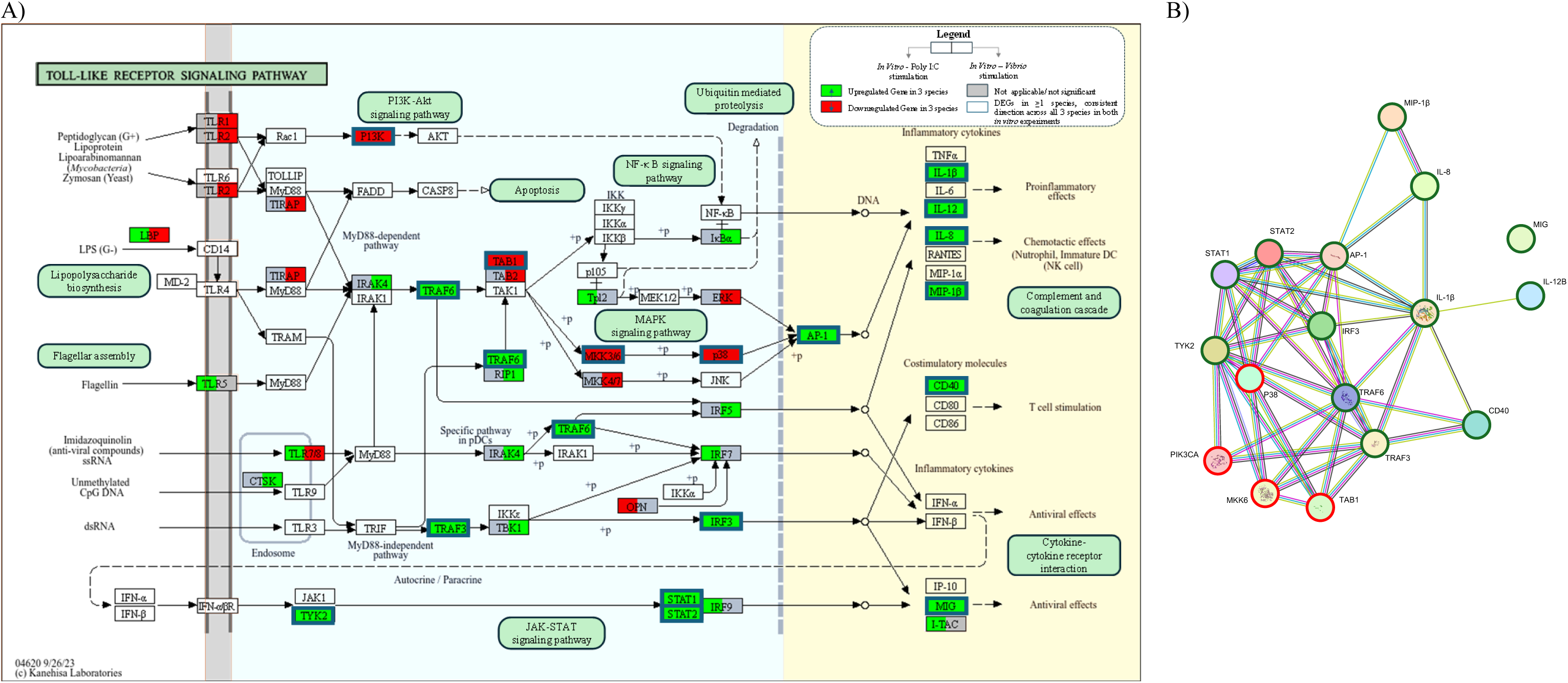
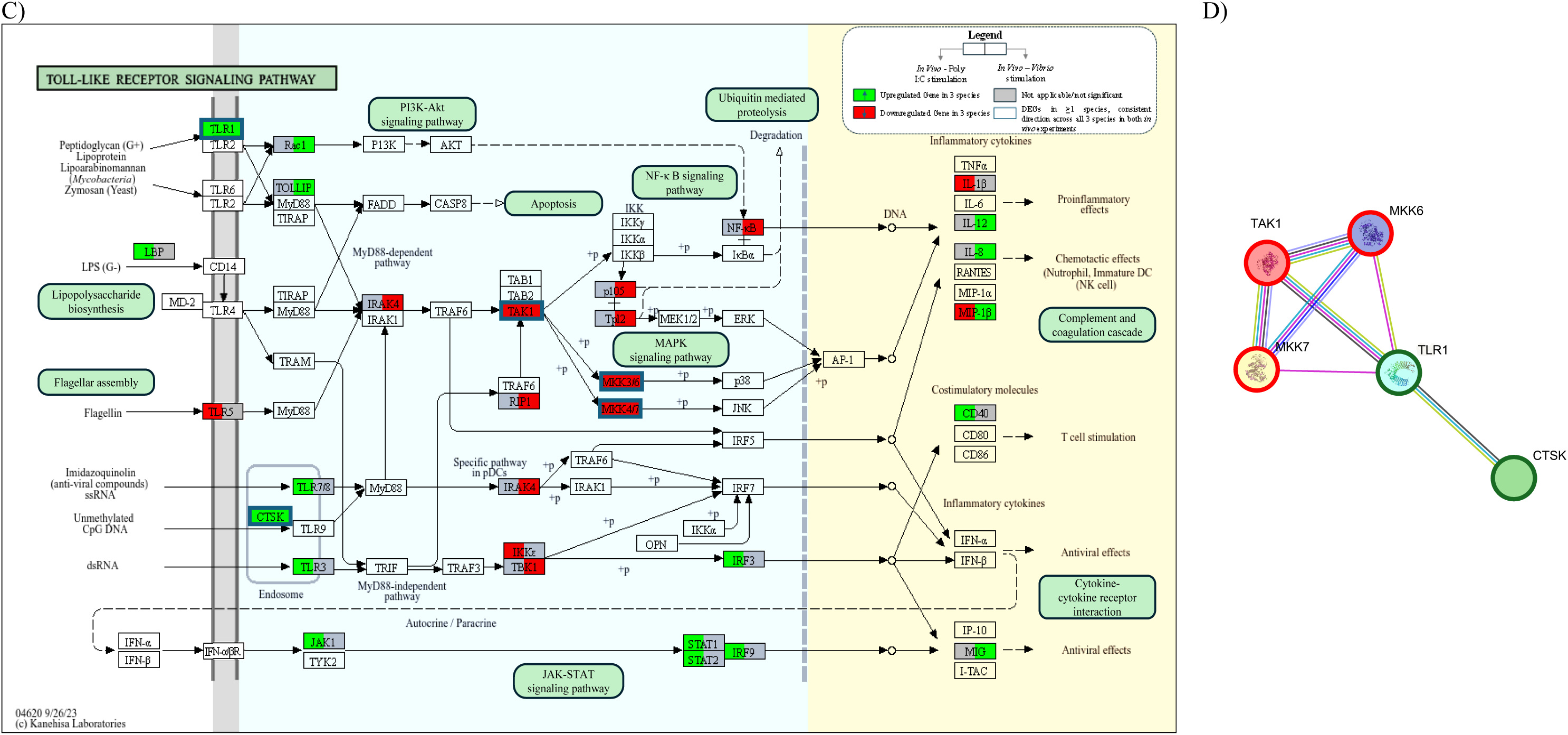
An overview of the toll-like receptor (TLR) signalling pathway under different experimental conditions. A and C represent the TLR pathway in *in vitro* and *in vivo* conditions, respectively. Genes upregulated in S*. maximus, S. aurata* and *D. labrax* are shown in green colour, while genes that are downregulated across the three species are represented in red; grey boxes indicate no consistent regulation across species or absence of differentially expressed genes in any species. In each gene, the left side corresponds to Poly I:C stimulation, and the right side to *Vibrio* stimulation. Dark blue squares represent genes that are regulated in the same direction in both experiments and include a statistically significant differentially expressed gene in at least one species (Table S10B and S10C, respectively). These figures are modified from KEGG map04620 [39]. Figures B and D show protein-protein interaction (PPI) network analysis of genes with the same regulation pattern across species and stimulations for *in vitro* and *in vivo* conditions, respectively. The network was constructed using the STRING database with a confidence interval threshold of 0.150, where nodes connected by green edges indicate upregulated genes and red edges downregulated genes. The lines connecting nodes represent different evidence of interactions.

**Fig. 5.**
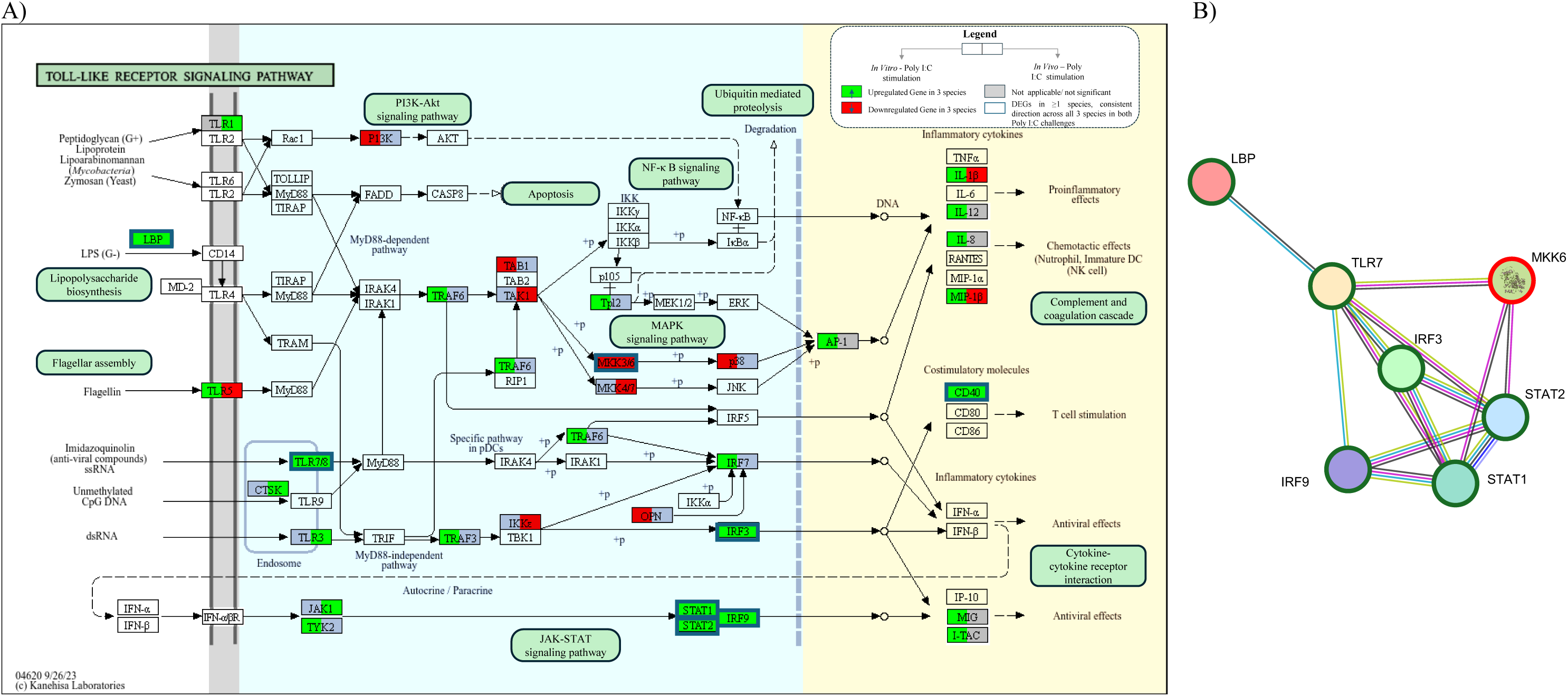
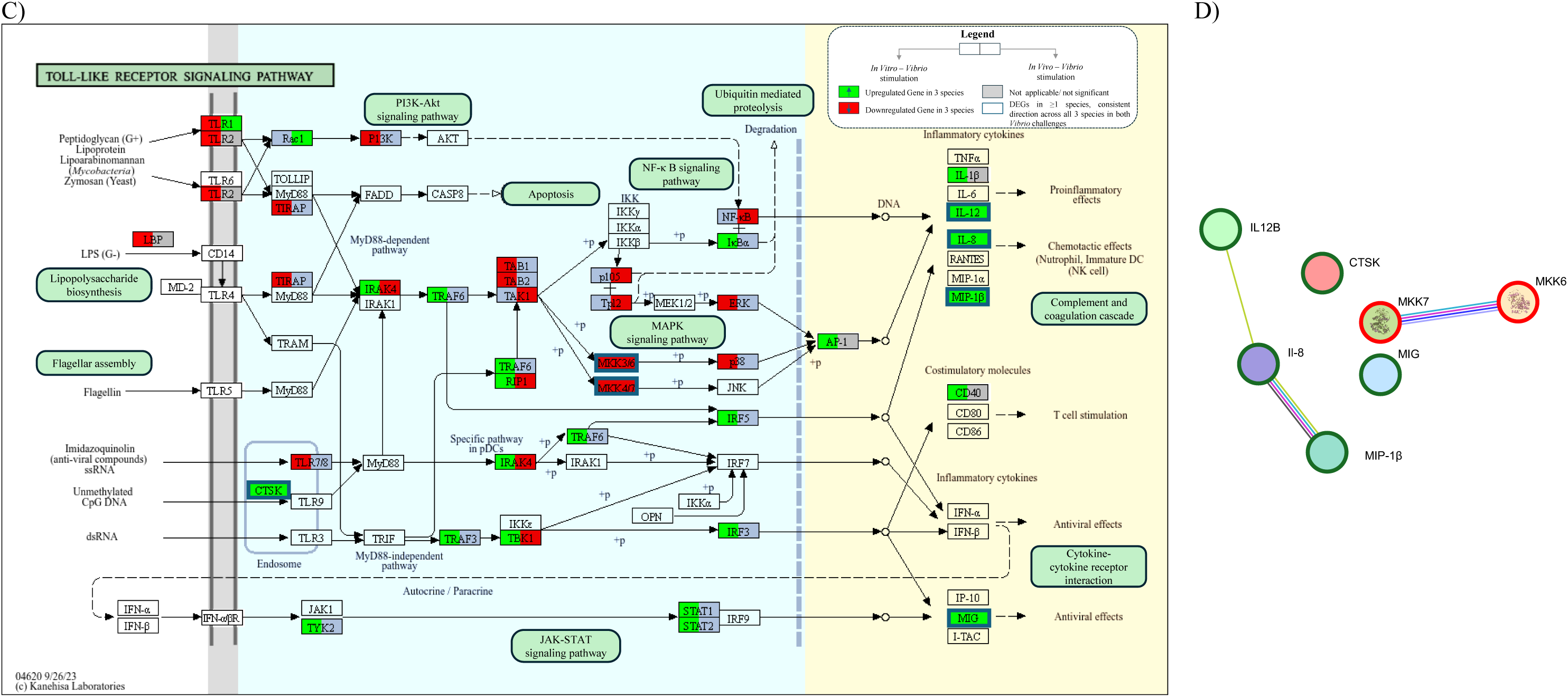
An overview of the toll-like receptor (TLR) signalling pathway under different experimental stimulants. A and C represent the TLR pathway under Poly I:C and *Vibrio* stimulants, respectively. Genes upregulated in S*. maximus, S. aurata* and *D. labrax* are shown in green colour, while genes that are downregulated across the three species are represented in red, grey boxes indicate no consistent regulation across species or absence of differentially expressed genes in any species. In each gene, the left side corresponds to the *in vitro*, and the right side to the *in vivo* condition. Dark blue squares represent genes that are regulated in the same direction in both experiments and include a statistically significant differentially expressed gene in at least one species (Table S10D and S10E, respectively). These figures are modified from KEGG map04620 [39] Figures B and D show protein-protein interaction (PPI) network analysis of genes with the same regulation pattern across species and conditions under Poly I:C and *Vibrio* stimulants, respectively. The network was constructed using the STRING database with a confidence interval threshold of 0.150, where nodes connected by green edges indicate upregulated genes and red edges downregulated genes. The lines connecting nodes represent different evidence of interactions.

## 4. DISCUSSION

Infectious diseases present a significant challenge to European aquaculture, impacting both animal welfare and the economic growth of the sector. Exploring the genomic basis of immune function and disease resistance in farmed fish has a high priority for improving health in aquaculture breeding and ensuring the industry’s sustainability. This study examined the immune responses of three commercially relevant fish species in Southern Europe, *S. maximus*, *S. aurata*, and *D. labrax*, chosen for their strong research foundation, evolutionary diversity, and their relevance for comparison with other aquaculture teleosts. In the current study, we investigated the impact of viral (Poly I:C) and bacterial (*V. anguillarum*) stimulations *in vitro* and *in vivo* on transcript levels in these species. This effort represents a first step towards elucidating the conserved cellular and organismal responses to pathogens in fish. Transcriptomic profiling was used to analyse gene expression changes, identify DEGs specific to each pathogen type and annotate their biological functions using the GO and KEGG databases.

### 4.1 Comparative transcriptomics: overview

To identify conserved and robust gene interaction networks, orthologous DEGs across the three species were compared at two levels: in *vitro* vs *in vivo* and Poly I:C vs *Vibrio*. The general observations suggested no common overlapping between *in vitro* and *in vivo* responses to *Vibrio*, whereas partial overlap was observed under Poly I:C stimulation, indicating better concordance for the viral challenge. This difficulty in finding overlap between the two conditions has also been documented in salmonids, where only ∼25% gene overlapping was found between *in vitro* and *in vivo* responses to Poly I:C [42]. In our multi-species comparison, the higher conservation of DEGs across the three species with viral stimuli suggests that a potent stimulant, such as Poly I:C, induces a more evolutionarily conserved response among teleosts, whereas the response to bacterial stimuli exhibits greater variability across experimental systems and species.

In addition, a conserved pattern was observed *in vitro* for all three species against both pathogens. A set of DEGs was identified in response to Poly I:C and *Vibrio* under *in vitro* conditions, but no overlapping DEGs were found *in vivo* between the two stimuli and all species. This highlights the complexity of the immune system when the DEGs identified within an entire organism are compared to a simplified cell culture setting. This complexity encompasses systemic immune interactions, tissue-specific and species-specific responses, and the microbiome’s role in bacterial and viral infections in a living organism. Beyond variations in immune responses among species to viral vs bacterial pathogens, there may also be differences in their response to both pathogens on different timescales. Our study focused on the 20-24 h post-challenge period, emphasizing the need for further studies at varying intervals to identify conserved responses between organisms.

### 4.2 Immune pathways enriched across fish species

To expand the scope of our analysis, we also examined genes and pathways that exhibited consistent regulatory patterns across all species and immunostimulants, focussing not only on DEGs between infection stages, but also exploring similar expression trends across species. This comparative approach revealed that the toll-like receptor (TLR) signalling pathway was significantly enriched across both conditions and challenges, emerging as a central hub of cross-species immune activation.

Shared genes among species revealed that the most enriched *in vitro* pathways were related to the main PRRs of the innate immune system. The PRRs family, which helps identify pathogens and initiate immune response, includes TLR, NLR (NOD-like receptor signalling pathway), RLR (RIG-I-like receptor signalling pathway), and CLR (C-type lectin receptor signalling pathway), among others [43]. These receptor families exhibit functional specialization in pathogen recognition: NLRs functioning as cytoplasmic sensors that identify intracellular bacterial components or stress signals; RLRs, such as RIG-I and MDA5, detect viral RNA within the cytoplasm and stimulate type I interferon production through MAVS signalling on mitochondria, which is vital for antiviral defence in both epithelial and immune cells; CLRs attach to carbohydrate structures found on fungi, bacteria and viruses, activating the Syk kinase and CARD9 pathway, which promotes cytokine production and antigen presentation [8].

In contrast to the PRR-dominated *in vitro* response, the *in vivo* one revealed enrichment of pathways associated with DNA replication and repair, which may reflect increased proliferation and activation of immune cells during the host response. This process plays a key role in restoring immune populations following exposure to pathogens [44], but also reflects elevated haematopoietic activity in general [45]. The divergence between *in vitro* and *in vivo* pathway enrichment patterns suggests that while direct pathogen recognition mechanisms dominate immediate cellular response, systemic immune activation involves broader cellular reprogramming and proliferative responses.

Analysis of stimulant-specific responses revealed distinct functional specialization of the fish immune system. Beyond the enriched TLR pathway, Poly I:C activated additional PRR-related pathways such as NLR or CLC, consistent with the multi-layered viral recognition mechanisms employed by innate immunity [46]. In contrast, *Vibrio* primarily enriched the cytokine-cytokine receptor interaction, emphasizing inflammatory responses and intercellular communication networks [47]. This pattern demonstrates that Poly I:C primarily triggers pathogen recognition cascades similar to those observed under *in vitro* conditions, while *Vibrio* exposure activates pathways focused on inflammatory signalling and immune cell coordination. These findings align with comparative studies of viral and bacterial stimulation in teleosts, which demonstrate pathogen-specific pathway enrichment patterns, supporting the concept that fish have evolved specialized immune recognition and response mechanisms tailored to distinct classes of pathogens [23,48].

#### 4.2.1 Toll-like receptors signalling pathway

All experiments enriched the toll-like receptor signalling pathway involving numerous genes conserved across all three species. The TLR signalling pathway involves TLRs, which are membrane-bound receptors located on immune cells such as dendritic cells and macrophages. These receptors recognize extracellular pathogens or their components (like LPS, flagellin, and viral RNA). Upon activation, they lead to NF-κB and IRF signalling, resulting in the production of cytokines and type I interferons [8]. The consistent enrichment of the TLR signalling pathway in all experiments, regardless of stimulus (viral vs bacterial) or experimental condition (*in vitro* vs *in vivo*), reveals a functional convergence toward evolutionarily conserved innate immunity mechanisms. As will be discussed later, this pathway includes numerous genes that belong to other pathways and are potentially conserved across all three species. Downstream signalling cascades converge toward common immune activation pathways [49,50], although specific ligands and primary TLR receptors vary between conditions, as observed in our study (e.g., conserved upregulation of TLR1 under *in vivo* conditions, and conservation of TLR7 and TLR3 genes in response to Poly I:C). Functional analysis of our data demonstrated this convergence through three fundamental aspects: i) the upregulation of genes associated with inflammatory cytokines or co-stimulatory molecules; ii) the negative regulation of the MAPK signalling pathway; and iii) the upregulation of the JAK-STAT pathway. The overexpression of genes linked to inflammatory cytokines or co-stimulatory molecules can result in excessive inflammation, hyperactivation of immune cells, and potential tissue damage or autoimmune-like symptoms, which are often observed in cytokine storms during severe viral infections [51]. Conversely, the downregulation of the MAPK signalling pathway, which plays a role in inflammation, cell survival, and proliferation, leads to reduced inflammatory signalling, decreased immune cell activation, and possibly a weakened response to bacterial infections or stress signals. This could serve as a protective mechanism against over-inflammation, or it might indicate immune evasion by pathogens [52]. Moreover, the upregulation of the JAK-STAT pathway, which transmits signals from cytokines like interferons to the nucleus to activate immune genes, plays a crucial role in antiviral defence. This upregulation increases the expression of antiviral genes, improves communication among immune cells, and strengthens the body’s ability to fight off viruses [53].

### 4.3 Key conserved genes

In addition, the comparative transcriptomic study revealed potentially conserved differentially expressed genes shared between *in vitro* and *in vivo* in responses to Poly I:C (Table S8), DEGs shared between Poly I:C and *Vibrio in vitro* stimulation (Table S9) and genes that demonstrated a uniform expression pattern in the four experimental challenges performed (DEGs and non-DEGs) (Fig. 3). This integrated approach allowed to investigate the most important genes involved in the three species’ response to viruses and bacteria, particularly those related to chemokines, interferon, antigen processing and presentation, cell signalling regulators and MAPK (for details, see Supplementary Discussion).

#### 4.3.1 Chemokine related genes

Chemokines, essential signalling molecules, play a fundamental role in modulating immune responses by recruiting immune cells to sites of infection and mediating communication between innate and adaptive immunity [54]. Notably, *ccl19* emerged as a primary gene, showing consistent upregulation both *in vitro* and *in vivo* in response to bacterial and viral pathogens across the three species studied (Table S9, Fig. 3). This gene is known to be involved in the recruitment of T cells and maturation of dendritic cells (DCs), thereby facilitating adaptive immune response. The TLR pathway identified in this study links *ccl19* upregulation with the production of pro-inflammatory cytokines (*il-12*, *il-1β*) and co-stimulatory molecules, consistent with its role in bridging innate and adaptive immunity during host defence responses in other species [55,56].

#### 4.3.2 Interferon-related genes

Interferons (IFNs) are a family of pleiotropic cytokines that represent a crucial part of the innate immune response against invading pathogens [57] by inducing hundreds of IFN-stimulated genes (ISGs) [58]. IFN regulatory factors (IRFs) are transcription factors regulating the expression of IFN and other related genes [59]. This study identified several conserved IFN-related genes in response to stimulators across the three species, including the viral sensor *ifih1*, IFN-induced genes such as *ifi35*, *ifi44,* and *isg15*, and regulators such as *irf3*, *etv7* and *uba7* (Table S8, S9). Infection with Poly I:C induced strong upregulation of *ifih1*, a helicase that detects viral dsRNA and activates type I IFN signalling cascades [60–62]. This process subsequently leads to the induction of *isg15*, a major interferon-stimulated gene involved in ISGylation [63]. The E1 enzyme *uba7*, along with other ligases, mediates this ISGylation by modulating the stability and activity of proteins during infection; *uba7* was also significantly upregulated, supporting its pivotal antiviral role [58,64]. Although knowledge regarding *ifi35*, *etv7,* and *uba7* in teleosts is still limited, *ifi35* is known to regulate pro-inflammatory cytokines and *ifn-β* [65], while *etv7* serves as a negative regulator that may prevent excessive inflammatory signalling [66]. Both genes emerged as conserved upregulated DEGs, indicating their role in controlling interferon responses in fish. Additionally, the conserved *ifi44*, which encodes an antiviral intracellular protein [67,68] and *irf3*, which is essential for mediating type I IFN and ISG expression [59,69], displayed increased expression in response to viral and bacterial stimuli, underlining their central roles in pathogen defence.

#### 4.3.3 Antigen processing and presentation-related genes

Upon the invasion of a pathogenic antigen into the cytoplasm, the initiation of antigen processing and presentation (APP) occurs [70]. We observed activation of this APP process in response to viral and bacterial stimuli, as evidenced by the upregulation of several DEGs in all species for both *in vitro* challenges: *psme1, psme2, psmb13a, tapbp.2, tap1, tap2a* and in Poly I:C in both conditions: *nlrc5* (Table S8, S9). Additionally, two other DEGs were upregulated in all four conditions: *psmb9a* and *erp44* (Fig 3). These comparative findings demonstrate that the molecules involved in APP are essential for the immune system, not only in cell models but also *in vivo*, showing their conservation in the three species studied and their conserved positive regulation in response to external infections across taxa.

#### 4.3.4 Cellular signalling regulators

In this study, we identified expression changes of several conserved cellular signalling regulators 20-24 h after stimulation (Tables S8, S9, Fig. 3). As they orchestrate immune responses, upregulation of these conserved genes aims to enhance immune activation, reflecting their roles in the release of antibacterial peptides (*bpifcl*) [71], immunomodulation (*lgals9l5*) [72,73], B cell receptor signalling (*igbp1*) [74], regulation of antiviral transcription and inflammatory genes (*stat1a*) [75,76] and feedback inhibition of cytokine signalling (*socs1b*) [77,78], respectively. Conversely, downregulation of *arrb2* and *itk* may reflect pathogen strategies to modulate surface receptor signalling [79] and, in the case of *itk*, also T cell activation for immune evasion [80].

#### 4.3.5 Mitogen-activated protein kinase

In the experiments, the *map2k6* gene was consistently downregulated across all conditions (Fig. 3). Examination of the TLR-MAPK pathway revealed that most genes also exhibited downregulation. Even though MAPK pathway upregulation would typically be expected due to its role in activating inflammatory cytokines and immune responses, the significant downregulation observed here in response to the experimental challenges suggests a distinct modulation of this pathway under these conditions. In addition, our study conducted at 20-24h could reflect the early activation of *map2k6* as an initial defence mechanism followed by negative feedback regulation [23,81]. Although this study did not directly investigate the underlying causes, it has been observed that *map2k6* expression can be significantly decreased in certain contexts, which could reflect an evolutionary strategy employed by pathogens to evade host immunity [25,82].

## 5. CONCLUSIONS

This study provides the first comprehensive comparative transcriptomic analysis for three acanthopterygian fish species, *S. maximus*, *S. aurata*, and *D. labrax*, in response to viral and bacterial infections, using both *in vitro* and *in vivo* conditions. The findings reveal that all three species exhibit remarkable changes in the expression of immune-related genes *in vitro*. In contrast, *in vivo*, they show an enhanced response related to DNA replication and repair. Additionally, viral and bacterial challenges elicited differentiated immune pathway activations, reflecting specialized host defence mechanisms. Moreover, a more conserved response was observed across the three species when exposed to the viral mimic than to the heat-killed gram-negative bacteria. Analysis of orthologous gene sets across the three species revealed conservation of genes involved in chemokines, interferons, antigen processing and presentation, cell signalling regulation, and MAPK pathways in response to external pathogens, with some of these genes reported for the first time as part of a shared response to Poly I:C or *Vibrio*. Our results demonstrate not only several of the deeply conserved pathways in immune responses among Acanthopterygian fish species, but also the use of the comparative method to uncover responses that may have been overlooked in single-species studies, e.g. due to differences between species in response time. This comprehensive approach offers valuable insights into immune responses across multiple species and conditions, while significantly contributing to the critical field of comparative immunology in aquaculture.

## Supporting information

Supplementary Result

Supplementary Discussion

## Credit Authorship contribution statement

R.R.V.: Conceptualization, methodology, software, formal analysis, investigation, data curation, visualization, writing - original draft, writing - review & editing; M.K.G.: Conceptualization, methodology, investigation, writing - review & editing; O.A.: methodology, data curation, investigation, resources, writing - review & editing; J.R.: methodology, data curation, resources, writing - review & editing; C.S.T.: funding acquisition, methodology, resources, project administration, writing - review & editing; S. F.: methodology, investigation, resources, writing - review & editing; R.F.: methodology, data curation, resources; L.B.: funding acquisition, methodology, resources, project administration, writing - review & editing; P.M.: funding acquisition, methodology, resources, project administration writing - review & editing; D.R.: methodology, investigation, supervision, resources, funding acquisition, project administration, writing - review & editing; H.J.M.: conceptualization, methodology, investigation, supervision, resources, funding acquisition, project administration, writing - review & editing.

## Acknowledgments

We acknowledge the technical support and informatic resources provided by the Centro de Supercomputación de Galicia (CESGA). We would also like to thank Pantelis Katharios, Tereza Manousaki, Ioannis Papadakis and Elena Sarropoulou for their help in gilthead sea bream sampling and preliminary analyses. Thank also certain AQUA-FAANG partners for fruitful discussions and protocol optimizations.

## Data availability

Raw RNA-seq datasets can be accessed through the ENA repository under the following accession number: turbot (PRJEB47933), seabream (PRJEB64880, PRJEB64877) and seabass (PRJEB52285, PRJEB52283). Detailed metadata for the samples and prepared libraries are available in Supplementary Tables S1 and S2, respectively. Detailed experimental protocols are publicly available in the FAANG repository (data.faang.org) and following the URLs facilitated in Supplementary Tables S1 and S2. Additional data will be made available on request. Supplementary data has been attached.

