## Supplementary Material & Methods for "Comparative transcriptomics of immune response to viral and bacterial stimuli in three acanthopterygian bony fish"

*Animals*

For this study, we collected samples from three different species: *Scophthalmus maximus, Sparus aurata* and *Dicentrarchus labrax.* Thirty turbot, twenty-six seabream and twenty-four seabass specimens were used for this study (Table S1). Turbot specimens were provided by Stolt Sea Farm S.A. (Ribeira, Spain), and housed in indoor tanks with recirculating seawater at the facilities of the Aquarium of the University of Santiago de Compostela (Spain) for 15 days of acclimation at 16ºC. Seabream specimens were obtained from Forkys S.A. Aquaculture in Sitia, Crete, whereas spawning took place in at the Aqualabs facilities of IMBBC, HCMR. The offspring was kept in indoor tanks with recirculating seawater maintained at 16ºC. The seabass specimens were provided from a commercial, NNV-free tested, broodstock from Valle Cà Zuliani Società Agricola srl (Pila di Porto Tolle, Rovigo, Italy); 2 years-old fish were then transferred to the Istituto Zooprofilattico Sperimentale delle Venezie (IZSVe, Legnaro, Padova, Italy). The fish were then kept in indoor tanks filled with artificial saltwater (30‰ salinity, temperature 20 ± 1°C, oxygen 6 ppm) and exposed to artificial photoperiod (14 h of light, 10 h of darkness).

All fish were deprived of food for 20 h (turbot), 24 h (seabream) and 48 h (seabass) before stimulations were performed. Eighteen fish were stimulated *in vivo* by intraperitoneal injection in each species, while the remaining were used for isolation of leukocytes for *in vitro* stimulation. Firstly, fish were anesthetized by bath (MS-222; 100 mg/L) and consequently euthanized with anaesthetic overdose (MS-222; 150 mg/L) prior to tissue collection. For turbot, all fish experiments were carried out in accordance with the Bioethics Committee of the University of Santiago de Compostela (body authorized according to R. D. 53/2013) and with the authorization of the Xunta de Galicia Regional Government. Meanwhile, all experimental procedures involving seabream were approved by the Bioethics Committee of the General Directorate of Rural Economy and Veterinary, Region of Crete, Greece (authorization no. 32356/09.02.2021), and conducted in full compliance with its ethical standards. For seabass, animal care was fully compliant with the prescriptions of the Directive 2010&63/EU of the European Parliament and of the Council, implemented at national level through the

D. Lgs 4 March 2014, n.26, and all experiments were authorized by the Italian Ministry of Health (auth. 641-2018-PR).
