## Supplementary Result for "Comparative transcriptomics of immune response to viral and bacterial stimuli in three acanthopterygian bony fish"

### Supplementary results

***RNA-seq***

*PCA of transcriptomic responses*

Figure S1 shows the PCA for the three species under investigation across *in vitro* and *in vivo* stimulation with mimics of viral (Poly I:C) and bacterial infection (*Vibrio*). In general, a clear separation can be observed between control samples and those exposed to stimulation. The *in vitro* Poly I:C stimulation revealed a notable clustering of samples, with *S. aurata* exhibiting the most pronounced separation along PC1 (64% of variance explained) and *D. labrax* the weakest (46%). However, separation along PC2 (15%) enabled discrimination between the control and Poly I:C-stimulated groups (Fig. S1A). Conversely, the *in vitro Vibrio* stimulation revealed a marked differentiation among species, with PC1 accounting for ~70% of the variance (Fig S1B). Regarding the *in vivo* Poly I:C stimulations, they also separated all species along PC1, ranging from 64% variance explained in *S. maximus* to 47% in *D. labrax*, in all three cases with no overlap between groups (Fig S1C). Finally, under the *in vivo Vibrio* stimulation, the lowest PC1 variance was present in *S. aurata* (33%), which, due to a higher PC2 explained variance (17%), allowed the separate clusterization of control and *Vibrio*-stimulated samples. Similarly, turbot and seabass showed higher PC2 values and PC1 separation of ~40% (Fig. S1D). Taken together, these PCA results demonstrate a clear clustering between the controls and the stimulated samples, which allows for robust genomic comparisons.

***Functional enrichment among differentially expressed genes***

Gene Ontology (GO) analysis identified significantly enriched biological process (BP) GO terms for DEGs in each treatment and species (Table S5). Due to the diversity of GO terms, these were summarised and visualised using the online tool REVIGO (Reduce + Visualise Gene Ontology) [1] to enable better interpretation, visualisation and clustering of similar GO terms, labelling each group with a representative and unique gene ontology term (Table S6). Due to the relevance of immune functions, immune/defence related GO categories were prioritised to discard irrelevant data, thus each cluster with direct association with immune response was selected. The REVIGO GO terms selected for each species/condition are shown in Table S6 and the bubble plot in Figure S4. The *in vitro*-Poly I:C study revealed that the species *D. labrax* exhibited the highest number of clusters (Fig. S4A) and GO terms (Table S6) associated with the immune response. Furthermore, it is noteworthy that the three species exhibit a number of common GO terms related to the immune response (e.g. GO:0002376, GO:0006955), response to external stimuli/pathogens (e.g. GO:0006952, GO:0009607, GO:0051707, GO:0043207), and biological interactions (GO:0044419) (Table S6, Fig. S4A). In contrast, the *in vitro-Vibrio* experiment discloses that turbot species exhibited a greater number of terms related to the immune response and to other stimuli than the other two species. Moreover, it should be noted that the terms "immune system process" and "immune response" were identified in all three species (Fig S4B, Table S6). Conversely, the *in vivo*-Poly I:C model showed a reduced number of clusters in *D. labrax*, while a greater degree of overlap was observed between turbot and seabream, encompassing immune system processes and immune responses (e.g. GO:0002376, GO:0006955, GO:0002682), response to stimuli (e.g. GO:0009607, GO:0006952, GO:0043207, GO:0009615) and interspecies interaction (GO:0044419) among other processes (Fig S4C Table S6). *In vivo-Vibrio* challenge demonstrated a greater number of enriched clusters in turbot species, indicating that the “stress response” is a structure shared by the three species. Between turbot and seabream, a higher number of overlapping GO terms was found related to general immune response (GO:0006955), response to specific pathogens (e.g. GO:0009615, GO:0009617), cell signalling and chemotaxis (GO:0006935), stimulus response (e.g. GO:0050896, GO:0009605), stress response (e.g. GO:0006950), and interspecies interaction (GO:0044419), among others (Fig S4D,Table S6). Associated differentially expressed associated genes (DEGs) were obtained from each of the selected GO terms and will be used in subsequent analyses for transcriptomic comparison studies between the three species.

***Turbot, seabream and seabass head kidney comparative transcriptome***

The list of immune response-associated DEGs was examined for orthology across the three species under study using BioMart and the Ensembl database.

*In Vitro*-Poly I:C stimulation resulted in a total of 503 DE-orthologs being differentially expressed in one, two or all three species. Among these, 31 DE-orthologs (6.16%) were shared across all three species (Table S7A, Fig. 2A). Turbot and seabream shared 27 DEGs (5.4%), seabream and seabass 18 DEGs (3.6%), and turbot and seabass only 2 DEGs (0.4%). Expression profiling revealed species-specific patterns, with turbot exhibiting higher expression of interferon-related genes, immune system regulators, signalling molecules, and cell surface patterns, while seabream and seabass showed elevated expression of genes linked to the proteasome, antigen processing and components of the immune system.

*In vitro-Vibrio* stimulation found a total of 1,472 DE-orthologs being differentially expressed in one, two or all three species. It was found 140 DEGs (9.5%) were shared among all three species (Table S7B, Fig. 2B). Pairwise overlap was the highest between turbot and seabream at 148 DEGs (9.5%), followed by turbot-seabass at 99 DEGs (6.7%) and seabream-seabass at 72 DEGs (4.9%). Expression profiling showed turbot again diverging from the other two species, with increased expression in genes associated with immune signalling pathways, including TNF signalling, cytokines, chemokines and interferon, as well as in genes related to immune cell receptors, signal transduction and other immune genes. In contrast, seabream and seabass exhibited higher expression levels than turbot in a distinct set of genes, such as cell signalling, immune response and cell regulation.

*In vivo-* Poly I:C response identified a total of 920 ortholog genes being differentially expressed in one, two or all three species, of which 22 DEGs (2.4%) were shared across species (Table S7C, Fig. 2C). The proportion of shared DEGs remained low, with only 17 (1.8%), 12 (1.3%) and 69 (7.5%) DEGs shared between turbot-seabream, seabream- seabass, and turbot-seabass, respectively. Although seabass displayed a more divergent expression profile, no discernible interspecies pattern emerged. Shared genes were mainly involved in both pro-inflammatory and anti-inflammatory signalling pathways and coordinating innate and adaptive immune functions.

*In vivo-Vibrio* challenge revealed the lowest level of overlap, with only 4 DEGs shared between the three species from 832 annotated ortholog genes being differentially expressed in one, two or all three species (Table S7D, Fig. 2D). Turbot and seabass showed the highest pairwise overlap with 91 DEGs (10.9%), followed by minimal overlap in other pairwise comparisons, with 12 DEGs (1.4%) between turbot and seabream, and with 14 DEGs (1.7%) between seabream and seabass. The expression pattern of shared genes anew highlighted a greater separation in seabass, with most genes showing higher expression in this species.

***Conserved immune responses across fish species***

*Candidate conserved genes expressed significantly in Poly I:C challenge*

A comparative analysis was conducted between the three fish species based on the previously identified and annotated orthologs in public databases. In order to understand how molecular mechanisms observed in cell cultures translate into physiological responses in living organisms, DEGs under both *in vitro* and *in vivo* conditions, in response to the same challenge and in the three species were identified. The aim was to determine which DEGs observed in cellular studies were also DEGs in living fish, thus establishing a correlation between experimental models and whole organisms. Our study identified the same DEGs in the cell cultures of all three species and in living fish against the Poly I:C stimulant, but not against *Vibrio* (Table S8). A total of 7 upregulated genes were identified in all species and in both *in vitro-in vivo* models, suggesting a similar behaviour in both cases. These upregulated genes in all species, in both systems and against the Poly I:C stimulus are bactericidal permeability-increasing protein fold containing family C, like (*bpifcl*), interferon-induced protein 35 (*ifi35*), interferon-induced protein 44-like (*ifi44*), interferon induced with helicase C domain 1 (*ifih1*), lectin, galactoside-binding, soluble, 9 (galectin 9)-like (*lgals9l*), NLR family CARD domain containing 5 (*nlrc5*) and signal transducer and activator of transcription 1a (*stat1a*).

*Candidate conserved genes expressed significantly in vitro condition*

In addition to the previous analysis, understanding molecular defence mechanisms against different pathogens is key to unravelling the molecular processes involved in immune responses. For this purpose, it was also needed to identify genes that exhibited significant differential expression *in vitro* in response to both *Poly I:C* and *Vibrio* in all species under study. This approach enabled the identification of genes that play a pivotal role in response to external pathogens and may therefore be functionally conserved [2]. Our comprehensive analysis uncovered a set of 15 DEGs in both *in vitro* experiments and in all three fish species studied (Table S9), but no shared DEGs were found for virus and bacterial mimic challenges in *in vivo* conditions. Remarkably, 13 out of these 15 genes showed a conserved pattern of regulation, with the same directional change in expression in response to both stimuli in all species. Among these genes, the *bpifcl* (BPI fold containing family C) gene was upregulated in response to Poly I:C stimulation in the three species, whereas it was downregulated in response to *Vibrio* across all species. Regarding the lymphocyte antigen 75 (*ly75*) gene, it was upregulated in *S. maximus* in both challenges, and in *S. aurata* and *D. labrax* it was upregulated in response to Poly I:C stimulation, while it was downregulated in response to *Vibrio*. The genes that were upregulated in all species and in both experimental studies are: chemokine (C-C motif) ligand 19 (*ccl19*), Interferon-induced protein 35 (*ifi35*), interferon regulatory factor 3 (*irf3*), interferon stimulated gene 15 (*isg15*), proteasome activator complex subunit 1 (*psme1*), proteasome activator complex subunit 2 (*psme2*), proteasome subunit beta 13a (*psmb13a*), suppressor of cytokine signalling 1b (*socs1b*), TAP binding protein (tapasin), tandem duplicate 2 (*tapbp2*), ETS variant transcription factor 7 (*etv7*), transporter 1, ATP-binding cassette, sub-family B (MDR/TAP) (*tap1*), transporter associated with antigen processing, subunit type a (*tap2a*) and ubiquitin-like modifier activating enzyme 7 (*uba7*).

*KEGG pathway enrichment analysis of immune responses across fish species*

In addition, genes that shared the same regulation (that were significantly differentially expressed in at least one species and showed the same direction of change in the other two species), defined as conserved regulation genes (CRGs). These CRGs were analysed separately across four different experimental combinations: *in vitro* studies (combining Poly I:C and *Vibrio* responses), *in vivo* studies (combining Poly I:C and *Vibrio* responses), Poly I:C stimulation (combining *in vitro* and *in vivo* responses), and *Vibrio* stimulation (combining *in vitro* and *in vivo* responses). The identified gene lists are presented in Table S10B-E. From the *in vitro* studies, a total of 140 CRGs were identified for the simultaneous interaction of Poly I:C and *Vibrio* (Table S10B), i*n vivo* studies found a total of 60 CRGs (Table S10C). A total of 54 CRGs were found under Poly I:C stimulation across *in vitro* and *in vivo* conditions (Table S10D) and a total of 90 CRGs were found under *Vibrio* stimulation (Table S10E).

Enrichment pathway analyses were conducted separately for each gene list using the Kyoto Encyclopaedia of Genes and Genomes (KEGG) database (see Table S11). The enriched KEGG associated with immune-related pathways are presented in Figure S5. *In vitro* studies revealed a total of nine pathways such as toll-like receptor signalling pathway (16 genes), NOD-like receptor signalling pathway (16 genes), cytokine-cytokine receptor interaction (17 genes), C-type lectin receptor signalling pathway (12 genes), RIG-I-like receptor singaling pathway (8 genes), necroptosis (10 genes), MAPK signaling pathway (15 genes), proteasome (5 genes) and cytosolic DNA-sensing pathway (5 genes) (Fig. S4A). In contrast, *in vivo* studies showed enriched KEGG for the following five pathways related to immune response, such as nucleotide excision repair (6 genes), DNA replication (4 genes), base excision repair (4), toll-like receptor signaling pathway (5 genes) and Mismatch repair (3 genes) (Fig. S4B). Under Poly I:C challenges four significant immune-related pathways were identified: toll-like receptor signalling pathway (7 genes), NOD-like receptor signaling pathway (6 genes), necroptosis (5 genes) and C-type lectin receptor signalling pathway (4 genes); meanwhile under *Vibrio* challenges it was found toll-like receptor signalling pathway (7 genes) and cytokine-cytokine receptor interaction (9 genes).

*Toll-like receptor signalling pathway*

A previous analysis demonstrated that the Toll-like receptor (TLR) signalling pathway is a common pathway of all experiments (Fig. S5). In the *in vitro* study, a total of 13 genes were found to be upregulated in all three species against Poly I:C and *Vibrio* (*traf6, traf3, irf3, tyk2, stat1, stat2, ap-1, il-1β, il-12, il-8, mip-1β, cd40, mig*), while 4 were downregulated against both stimuli (*pik3ca, tab1, mkk6, p38*). In addition, two pathways were found with a higher percentage of downregulated genes, PI3K-Akt signalling pathway (such as *tlr1, tlr2, pik3ca*), and the MAPK signalling pathway (with *tlr2, tirap, tab1, tab2, mkk6, mkk7, erk, p38*). On the contrary, there is an upregulation of the JAK-STAT signalling pathway (with *tyk2, stat1, stat2, irf9*), and in the nucleus most of the genes are related to inflammatory cytokines and molecular co-stimulators (such as *ap-1, il-1β, il-12, il-8, mip-1β, cd40, mig, i-tac*) (Fig. 4A). The analysis of the internodal relationships between genes expressed in the same direction in both experiments revealed some connectivity patterns (Fig. 4B). The central genes show interconnections with a distinct separation between those that are upregulated and those that are downregulated. These interconnections, as reported by STRING, represent predicted functional associations supported by multiple sources of evidence (experimental data, curated databases, co-expression and text-mining). A stronger link is observed between anti-inflammatory cytokinin and co-stimulatory molecules, which form a cohesive group. Similarly, genes involved in the JAK-STAT pathway tended to cluster together, indicating a coordinated regulation of the signalling pathway. The molecules *irf3*, *traf6* and *traf3* stand out as central nodes, acting as convergence points for multiple signalling pathways.

The TLR signalling pathway map of the genes found in *in vivo* study is shown in Fig. 4C. A lower number of genes were found to be regulated in the same direction against both *in vivo* stimuli (Table S10C). Two genes (*tlr1, ctsk*) were found to be upregulated in all three species against Poly I:C and *Vibrio* in living organisms, while three genes (*tak1, mkk6, mkk7*) were downregulated against both stimulants in all species. The TLR5-NFKB signalling pathway exhibited a significant downregulation of its genes, including *tlr5, irak4, tak1, p105, tpl2,* and *nf-kβ*. Additionally, genes belonging to the MAPK signalling pathway, such as *irak4, tak1, p105, tpl2, rip1, mkk6,* and *mkk7*, also demonstrated downregulation. Alternatively, the analysis identified the presence of upregulated genes belonging to the endosome (*ctsk, tlr7, tlr3*) and the JAK-STAT signalling pathway (*jak1, stat1, stat2, irf9*). Fewer genes were found to be upregulated to express inflammatory and co-stimulatory cytokines than in cells *in vitro*, but *il-12, il-8, cd40, mig, mip-1β* were upregulated, while *il-1β* and *mip-1β* genes were downregulated against Poly I:C. Furthermore, a network analysis was conducted to examine the interrelationships between the genes that were expressed in the same direction in both experiments (Fig. 4D). The results indicated that the downregulated genes exhibited a higher degree of interaction with each other, whereas the upregulated genes demonstrated a greater interconnectivity in *tlr1*, which served as a pivotal node between the two groups.

A detailed analysis of the TLR signalling pathway revealed a heterogenous patterns under Poly I:C stimulation (Fig. 5A, Table S10D). Seven genes (*lbp, tlr7, irf3, irf9, stat1, statt2* and *cd40*) were upregulated in all three species against Poly I:C, both *in vitro* and *in vivo*, while *mkk6* was a CRG downregulated against this stimulant and both conditions. Moreover, an upregulation of key pattern recognition receptors (*tlr1, tlr3, tlr7*), adaptor proteins (*lbp, ctsk, traf3, traf6*), and downstream effectors (*irf3, irf7*) was shown, while CRGs involved in the PI3K-Akt signalling cascade (*pik3ca*) and MAPK signalling pathway (*tab1, tak1, mkk6, mkk7, p38*) were downregulated. The pathway analysis revealed stronger activation of the JAK-STAT pathways components and IRF-mediated antiviral responses in both conditions (*stat1, stat2, irf9, irf7*). These genes, along with key inflammatory cytokines (*mig, i-tac*), highlighted *traf3*, *traf6*, and *irf3* as central regulatory nodes connecting multiple signalling cascades, such as the expression of *cd40*. Notably, most inflammatory cytokines were upregulated under *in vitro* condition (*il-1b, il-12, il-8, mip-1b*), but not *in vivo*-Poly I:C. Furthermore, a network analysis was conducted to examine the interrelationships between the genes that were expressed in the same direction in cells and living organisms under Poly I:C stimulation (Fig. 5B). The results indicated that *irf3* was a central node with a higher degree of interaction between different genes.

Regarding the TLR signalling pathway associated with *Vibrio* stimulation (Fig 5C), the analysis revealed that five CGRs were upregulated (*cstk, il-12, il-8, mip-1β* and *mig*) and two downregulated (*mkk6* and *mkk7*). Noteworthily, this pathway shows that key negative regulators and components of PI3K-Akt and MAPK cascades (e.g. *pi3k, tirap, nf-kb, tab1, tab2, p105, tpl2, erk, mkk6, mkk7, p38*) are predominantly downregulated. Interestingly, although the upregulation of the JAK-STAT pathway components (*tyk2, stat1, stat2*) was apparent, this pattern appeared less conserved than in other experimental conditions. Moreover, *Vibrio* stimulation triggered a pronounced cellular response mediated by inflammatory cytokines and co-stimulatory molecules. Many of the previously mentioned CGRs, as well as *il-1β* and *cd40*, were upregulated, underscoring the central role of pro-inflammatory and immune-modulating signals in the antibacterial response. In addition, network analysis revealed no clear interactions among different CGRs under *Vibrio* stimulation, except between *mkk6* and *mkk7* genes (Fig 5D).
