## Supplementary Discussion for "Comparative transcriptomics of immune response to viral and bacterial stimuli in three acanthopterygian bony fish"

*Chemokine related genes*

Chemokines play a crucial role in the immune response, regulating cell migration and leukocyte activation at infected sites, and establishing a connection between innate and adaptive immunity [1–4]. In teleost, chemokines were also found to have similar functions to those in mammals [1,5–7]. In particular, CC motif chemokine 19 (*ccl19*) is a homeostatic cytokine that belongs to the CC chemokine subfamily that plays a role in signalling proteins [8,9]. It is noteworthy that in our research, one of the most significant immune genes in our species is indeed *ccl19* (Table S9, Fig. 3), although other coding genes of chemokines (*cxcl9, ccl4*) and receptors (*cxcr3, cxcr2*) were upregulated in the three species in response to both viral and bacterial challenges *in vitro* (Table S10B). This is reflected in our KEGG pathway analysis with a statistically significant enrichment of the cytokine-cytokine receptor interaction pathway. *Ccl19* is responsible for regulating the recruitment of CCR7-expressing T cells, as well as the homeostatic trafficking of lymphocytes and dendritic cells under physiological conditions [10,11]. In our study, the *ccl19* gene is the only gene that appears statistically upregulated *in vitro* against virus and bacterial mimics in all three species (Table S9) and, at the same time, upregulated in all fish species, *in vivo*, against both mimics (Fig. 3). These findings are corroborated by numerous studies in fish, which have demonstrated increased mRNA expression following viral infections *in vivo* and *in vitro* [12–14] or its antibacterial role of *ccl19* by their upregulation against bacteria [6,15–18]. Noteworthy, *ccl19* is hypothesized to enhance T cell proliferation through the maturation of dendritic cells (DCs), as well as by increasing co-stimulatory molecules and the release of pro-inflammatory cytokines (*il-12, tnfα,* and *il-1β*) [11,19]. It was demonstrated in our TLR pathway map, where it was an upregulation of inflammatory cytokines (*il-12, il-1β*) and co-stimulatory molecules in all species during *in vitro* and under *Vibrio* exposure. The consistent upregulation of *ccl19* and other chemokines in our study underscores its potential significance in the fish immune response to diverse pathogens, suggesting it may play a crucial role in enhancing host defence mechanisms across multiple fish species, as previously observed in turbot in the *in vivo* study by Chen et al. (2013) [20].

*Interferon related genes*

Interferons (IFNs) constitute a family of pleiotropic cytokines recognized for providing a robust line of defence against invading pathogens [21]. These interferons can induce the expression of hundreds of interferon stimulated genes (ISGs), whose products control responses to pathogens [22] and, in turn, interferons and ISGs are regulated by a small family of transcription factors called Interferon Regulatory Factors (IRFs ) [23]. In our study we found several possibly conserved DEGs in the three species: viral sensors as *ifih1*, interferon-induced genes such as *ifi35, ifi44, isg15*, and regulators such as *irf3, etv7* and *uba7*.

Infection with Poly I:C triggered upregulation of a potentially conserved gene in both cell lines and the organism, the *ifih1* gene. This gene is expected to be upregulated as it is known to be involved in an antiviral response in fish [24–26]. *Ifih1* is a viral sensor gene encoding melanoma differentiation associated protein 5, a helicase enzyme belonging to RLR, which detects viral dsRNA and activates a signalling cascade leading to IFN I production [24]. These interferons then induce the expression of ISGs such as those found in this study *ifi35* (also known as *ifp35*), *ifi44* and *isg15* [27–29]. Two interferon-induced proteins, *ifi35* and *ifi44*, are identified as playing a significant role in the innate immune response against pathogens in several species. *Ifi35* was observed upregulated both *in vitro* and *in vivo* (Table S8) and in Poly I:C and *Vibrio* (Table S9). This ISG is induced by IFN and translocated from the cytoplasm to the nucleus, participating in the regulation of pro-inflammatory cytokines and *ifn-β* through multiple signalling pathways [30]. Although there are limited reports on *ifp35* in teleosts, and particularly in our species of interests, its antiviral and antibacterial functions have been reported [31–34]. Gao et al. (2022) [30] observed a significant increase in *ifi35* expression in *Epinephelus coioides* spleen cells following exposure to red spotted grouper necrosis virus, Poly I:C or LPS, which is consistent with our results showing upregulation in all scenarios. The consistent upregulation and the scarcity of studies in our species opens a window to delve deeper into the role of *ifi35* in response to various pathogens, suggesting that it may be a conserved gene across species, playing a crucial role in pathogen defence mechanisms. Similarly, *ifi44* was upregulated both *in vitro* and *in vivo* in response to Poly I:C. In teleosts, this gene has been found to be rapidly induced during viral infections [25,34,35], as it encodes an intracellular protein with anti-proliferative activity [29,36].

In addition to these genes, one was found to be the most highly expressed ISGs upon viral or bacterial infections; the interferon-stimulated gene 15 [37]. *Isg15* is an ubiquitin-like protein that can conjugate to cellular proteins through the process of ISGylation [38], affecting their localisation, stability or activity, and modifying host and pathogens proteins. Similarly to ubiquitination, ISGylation is a process that is mediated by the consecutive action of E1 (*uba7*), E2 (*ube2l6*) and E3 ligases, which bind *isg15* to the target proteins [22]. Furthermore, it has the function of a cytokine, inducing IFN-γ in T cells, stimulating natural killer cell proliferation and acting as a chemotactic factor for neutrophils [39,40]. The present study revealed the overexpression of *isg15 in vitro* for both pathogens and in all species under investigation (Table S9). The upregulation of fish *isg15* genes in response to viral infections has been demonstrated in a multitude of fish species [41–45] as well as in response to bacterial infections [42,46,47]. Furthermore, *isg15* has been identified as a protein that is expressed in response to pathogen entry in our species: In seabream, induction of *isg15* was demonstrated through Poly I:C and viral infections *in vitro*, thereby illustrating its antiviral defence system in this species [48]; in European seabass, it was found that *isg15* was upregulated at an early stage in the head kidney in response to Poly I:C and another virus [49]; in turbot, induction of *isg15* was observed following Poly I:C administration under *in vivo* conditions [50]. These findings are in accordance with our results suggesting that *isg15* may be a conserved gene with antiviral and antibacterial functions, playing a pivotal role in combating external pathogens. Interestingly, we observed that the gene encoding the ISGylation-activating enzyme *uba7* (previously denoted *ube1l*) was also significantly upregulated against viral and bacterial mimics *in vitro*. As mentioned above, this E1 enzyme is part of the ISGylation process, facilitating the covalent attachment of *isg15* to target proteins. In addition to the antiviral/antibacterial function previously demonstrated with *isg15*, studies in mice have revealed that *uba7* also plays a crucial role in antiviral defence, as its mutation has been shown to increase susceptibility to death following viral infection [51]. Despite its importance, there are few studies examining *uba7* expression in response to infections. For this reason, our study is particularly relevant in demonstrating the upregulation of *uba7* in three fish species in response to viral and bacterial infections.

As discussed above, during responses to pathogens it is not only a matter of activating all the response mechanisms, but also to fine tune them to avoid exacerbated responses that ultimately cause damage to cells. Interestingly, two regulators emerged in response to the pathogens in our study, *etv7* and *irf3*. *Etv7* is a conserved DEG in all three species in our study, displaying upregulation *in vitro* in response to both bacterial and viral stimuli. Recent research has revealed that *etv7* functions as a negative regulator of the type I interferon response. In particular, it acts as a transcription factor that negatively regulates ISGs, including *isg15*, during the early stages to prevent accumulation. An increase in *etv7* levels has been demonstrated to diminish the capacity of cells to impede viral infection. Nevertheless, this downregulation is vital to avert excessive inflammatory signalling and to prevent prolonged and extreme immune responses [52]. A limited number of studies that have identified *etv7* as a factor in the immune response to viral and bacterial pathogens in fish. Our study represents the only comparative investigation to provide evidence that *etv7* may act as a conserved gene in the immune response of fish. Given that ISG15 was also upregulated, *etv7* may play a pivotal role in fish, specifically in the control of interferon responses. Controlling ISG levels is therefore crucial, as insufficient levels can result in uncontrolled viral spread, while excessive activation can lead to interferon-related pathologies.

Interferon regulatory factors (IRFs) constitute a small family of transcription factors that regulate anti-infective pathway signals in the cell. This function is achieved through binding to regulatory elements that induce the expression of interferons, ISGs, pro-inflammatory genes and immune-related genes. As a result, IRFs are critical mediators of innate immune signalling [53–56]. Our findings indicate that *irf3* is conserved across the species examined in the *in vitro* study, exhibiting a response to both viruses and bacteria. It is relevant to note that *irf3* is a gene that may be conserved across all three species under study and is involved in antiviral defence, as evidenced by our TLR map under *Poly I:C* stimulus. A plethora of studies have demonstrated that the expression of *irf3* is upregulated in fish infected with Poly I:C or viruses [12,23,54,57–60]. Furthermore, *irf3* has been found to possess a pivotal role in antibacterial processes, as evidenced by its upregulation *in vitro*, and corroborated by numerous investigations in the scientific literature. This dual role in the immune response, both antiviral and antibacterial, highlights the significance of *irf3* in the fish immune system [59,61–63].

Collectively, these findings highlight the cross-species consistency and underscore the evolutionary conservation of the interferon-related genes *ifih1, irf3, isg15, uba7, etv7, ifi44* and *ifi35*, emphasising a central role for *isg15* and reinforcing its pivotal role in ISGylation as a response to pathogens. Furthermore, it is worth noting the fact that the upregulated *uba7, etv7* and *ifi35* are genes for which there is a lack of information in the literature regarding our species and their role in pathogen defence, opening possible avenues for future studies due to their potentially conserved role in pathogenic diseases. Further research into the specific functions and regulatory mechanisms of *ifi35* in teleosts could provide valuable insights into the evolution of innate immune responses and identify targets for enhancing disease resistance in aquaculture species.

*Antigen processing and presentation related genes*

Upon the invasion of pathogens into the cytoplasm, the process of antigen processing and presentation (APP) is initiated. This process involves the transformation of an antigenic protein into peptides, their loading and subsequent transportation to the cell surface by major histocompatibility complex (MHC) proteins, and the recognition of these peptides by CD8+ T cells [64]. The process of proteolysis in the cytosol is carried out by the immunoproteasome, which is stimulated by IFNγ. In mammals, this process is initiated by the replacement of the proteasome components *psmb5*, *6* and *7* by *psmb 8, 9* and *10*. In addition to the PSMB8 subunit, unique subunits of *psmb9*, termed *psmb12*, have been identified in teleosts, along with an additional subunit of *psmb10*, designated *psmb13* [65]. Moreover, two additional subunits, designated *psme1* and *psme2*, are implicated in the regulation of the immunoproteasome, although their role remains uncertain [66]. Subsequently, the peptides produced enhance the binding of the *tap1/tap2* channel transporters. In the endoplasmic reticulum (ER), an empty molecule of MHCI is stabilised by *calr, pdi,* and *erp57*, and is bound by the *tap* transporter via tapasin (*tapbp*) [67]. Once transported into the ER, the peptides are trimmed by *erap1/2*, where it has been shown that *erp44*, among other functions, can negatively regulate *erap1* to limit peptide trimming and regulate antigen presentation [68]. Lastly, peptides conjugated to MHC I molecules are transported via the Golgi apparatus to the plasma membrane at the for recognition by CD8+ T cells [69]. Part of MHC class I genes may be regulated by *nlrc5*, which associates and transactivates their promoters [70]. In our study, we observed a clear activation of this APP process to cope with viruses and bacteria, as we found DEGs upregulated in all species *in vitro* for both challenges: *psme1, psme2, psmb13a, tapbp, tap1, tap2a,* and in Poly I:C stimulation in both models: *nlrc5* (Table S8, S9). Moreover, *psmb9a* and *erp44* were upregulated in all four experiments: (Fig 3).

Some studies have examined the post-infection upregulation of immunoproteasome-related genes. *Psme1* and *psme2* have been found to be upregulated in response to infection with gram-negative bacteria in yellow catfish [17], grass carp [71], Japanese flounder [72,73], rainbow trout [74], as well as in zebrafish with upregulated protein expression infected with viruses [75]. The *psmb9* gene was also shown to exhibit increased expression in Atlantic salmon cells in response to Poly I:C and viral infection [76]. No literature was identified that specifically addressed the role of *psmb13a* in antiviral or antibacterial responses, and lack of information highlights the need for future research to explore their role in immunity and their biological relevance. *Tap1, tap2,* and *tapbpa* genes demonstrate a remarkable immune response across multiple fish species. In Japanese flounder and turbot, *tap1* is upregulated against gram-negative bacteria [73,77,78], while in threadfin fish, there shows activation of *tap1/2* against gram-positive bacteria [79]. Rock bream experiences an upregulation of *tap1/2* when exposed to viruses in red blood cells [80], while Sockeye salmon induces these genes with poly I:C [81]. For *tapbpa*, increased expression is observed in Atlantic salmon due to viral infections [76], in *Epinephelus coioides* and large yellow croaker by gram-negative bacteria [80,82], and by LPS in rainbow trout [83]. Meanwhile, *tapbp* was increased post-infection with gram-positive bacteria in threadfin fish [79]. Conversely, Liyanage et al. (2019) [68] observed an increase in *erp44* expression in the kidney after the induction of inflammation with LPS and with *S.iniae* at 24 hours, and with *E. tarda* and Poly I:C stimulants at 72 hours, in the big-belly seahorse. They postulated that this may be important in the regulation of APP level. Simultaneously, *nlrc5* has been found to be induced by Poly I:C/virus in teleost [13,53,84,85], as it helps to regulate both constitutive and inducible expression of MHC class I genes [86]. Our studies provide the first comparative analysis demonstrating the conservation of these molecules involved in APP. They appear not only in cell line studies but also in *in vivo* studies, showing their conserved nature in the three species studied and their positive regulation in response to external infections.

*Cellular signalling regulators*

Genes that are involved in cell signalling regulation are key for successful cell defence processes. In the studied fish, we found five upregulated (*bpifcl, lgals9l5, igbp, stat1a* and *socs1b*) and two downregulated conserved genes (*arrb2* and *itk*).

*Bpifcl* is a member of a gene family that encodes for antibacterial peptides released by neutrophils and that bind to lipopolysaccharides [87], although the function of *bpifcl* is not very clear. In zebrafish, *bpifcl* has been shown to regulate the expression of Kisspeptin, a neuropeptide that is affected by inflammation in the brain [88], while in our study it was observed that this gene was upregulated both *in vitro* and *in vivo* against Poly I:C, whereas it was downregulated against *Vibrio*. Our findings align with those of Aramburu et al. (2025) [89], suggesting that this gene could play a role in host defence functions.

*Lgals9l5* (also known as galectin-9) is a tandem receptor-type member of the galectin family that binds to carbohydrates with B-galactosidases [90]. Galectin modulates a number of immune responses through its binding to the T-cell immunoglobulin and Tim-3 ligand [91], reducing T-helper 17 and TH1 activity [92] and regulating cellular NK function [93], among other functions. In teleosts, although studies are more limited, antibacterial [94] or antiviral functions have been identified in yellow croaker species [95] and in *Planiliza haematocheilus* [96], whose expression increased significantly after exposure to Poly I:C, suggesting a fundamental role in the innate immune response. This last study has demonstrated the capacity of *lgals9* to neutralise viruses by binding to viral glycoproteins, inhibit the binding of virions to host cells by activating autophagy-related proteins, and even promote the degradation of viral particles by autophagy mechanisms, consequently inhibiting virus replication [96]. Given the scarce data regarding *lgals9l5* involvement in response to viruses in our species, further research is required to elucidate the precise mechanisms and to investigate its potential in the development of disease resistance strategies for these aquaculture species.

*Igbp1*, also known as alpha4, is a key signalling molecule in the immune system that associates with the Ig-α surface receptor and is involved in the signalling of the B cell receptor complex [97]. *Igbp1* acts as a regulatory subunit of the PP2A phosphatase, directing its activity towards PI3K and AKT. PP2A controls several cell signalling pathways, including apoptosis, proliferation, cytoskeletal organisation and cell migration [98]. In response to cytokine activation of GqPCR, *igbp1* facilitates the switch of pp2a from pi3k (forming the PP2Ac-IGBP1 dimer) to *akt* (establishing the PP2Ac-IGBP1-PP2Aa trimer), leading to inactivation of the PI3K/AKT pathway and promoting apoptosis [99]. It was found that *pi3k* was downregulated (see TLR maps), which could be a cellular response to inhibit viral replication, as many viruses use the PI3K/AKT pathway for replication [100]. However, our *igbp1* was found to be upregulated in all challenges, either by promoting the formation of the PP2Ac-IGBP1-PP2Aa complex (promoting dephosphorylation to inactivate *akt* and trigger apoptosis as a defence mechanism against viral infection), or the virus could be manipulating these pathways to its advantage (altering the balance between cell survival and apoptosis). The conservation of *igbp1* and its role in the regulation of immune processes could be reflected in our study. However, due to the limited number of studies on this gene in fish, further research is needed to determine its fundamental role in the immune response to pathogens.

As mentioned above, cytokines induce an intracellular signalling cascade that triggers inflammatory responses. This intracellular signalling process is controlled by the JAK-STAT pathway [101], hence the differentially expressed *stat1a* gene was found both *in vitro* and *in vivo* (Table S8) and the entire pathway is upregulated in different environments (see TLR maps). Activation of JAKs causes phosphorylation and activation of STATs, which translocate to the nucleus and bind to DNA regulatory sites to control antiviral transcription and inflammatory genes [102,103].

However, immune system cells induce repressor genes to inhibit the overexpression of inflammatory cytokines in order to moderate this response and maintain protective responses [104]. One of them are the intracellular proteins called SOCS (suppressors of cytokinin signalling) [105,106].

In the present study, it was observed that *socs1b* was significantly upregulated in all three species in the *in vitro* study against viral mimetics and bacteria (Table S9). It is in accordance with viral [53,58,104,105,107–110] and bacterial infections studies [17,73,108,111,112] in teleost. SOCS1b proteins act as inhibitory regulators of the JAK/STAT pathway at post-translational level [108,113] through the binding of SH2 domain of SOCS1 to the phosphorylated tyrosine of JAK proteins, inhibiting their activity. This is via ubiquitin-mediated degradation, a process of proteolysis involving ubiquitin ligases recruited to the 'SOCS box' motif of SOCS proteins [114]. We hypothesise that our upregulation of *socs1b* expression is an attempt to regulate the overexpression of our JAK/STAT pathway, which is activated by both Poly-I:C and *Vibrio*-infected fish. Due to the enhanced expression of the *jak* and *stat1/2* genes, the expression of *socs1* will be elevated in an effort to suppress the activity of JAK proteins and, consequently, the entire downstream cascade that they regulate, with the objective of modulating the inflammatory response and preventing cellular damage.

β-arrestin 2 (*arrb2*) is a multifunctional adaptor that regulates the signalling of various cell surface receptors and plays a complex role in the immune response [115]. Its effects may be either pro- or anti-inflammatory, depending on the context (reviewed in Freedman and Shenoy (2018) [116]). It was found that in response to viral stimuli, *arrb2* expression is suppressed by acetylation and ubiquitination, which is considered a coevolutionary immune evasion strategy [115]. A recent study in lepidopteran insects demonstrated that the overexpression of *arrb2* resulted in an increase in viral replication, whereas the silencing of the gene led to a reduction in viral replication [117]. Given the limited number of studies on *arrb2* in our three species, it remains uncertain whether the observed reduction in expression levels is a strategy of the pathogen to suppress the immune response or a host response to prevent pathogen replication. Due to its role in regulating *arrb2*, it could emerge as a potentially novel target for the development of antiviral and antibacterial strategies. However, the lack of species-specific studies underlines the need for further research to elucidate the exact role of *arrb2* in each pathogen-host context.

The downregulation of the *itk* gene observed in our study has important implications for the immune response and susceptibility to infection. *Itk* plays a crucial role in T cell receptor (TCR) signalling and is essential for T cell development, activation, and function, as well as for the integration of TCR downstream pathways and cytoskeletal reorganisation [118–120]. Despite its importance in the immune response to viruses [34] or bacteria [121], recent studies have shown that *itk* inhibition or downregulation can have beneficial effects in certain contexts. For example, *itk* inhibition blocks HIV replication at multiple stages of the viral cycle, including entry, transcription, and virion assembly/release [122]. A recent study also found that *itk* was downregulated against *Vibrio*, suggesting suppression of T-cell activation upon infection [123]. The findings of our study indicate that *itk* is downregulated, in conjunction with the observed downregulation of PI3K in TLR maps, suggesting the existence of a negative signalling cascade that could potentially modulate the immune response to viruses and bacteria, as previously demonstrated by Wang et al. (2015) [124]. This downregulation may represent an evasion mechanism employed by various types of infectious agents, both *in vitro* and *in vivo*, through alterations in cellular activation and function.

*Mitogen-activated protein kinase*

In the experiments performed, the *map2k6* gene was found to be consistently downregulated (Fig. 3). Upon examination of the TLR maps within the MAPK pathway, it was observed that most of the genes involved where downregulated (detailed in Figures 4 and 5). The mitogen-activated protein kinase (MAPK) signalling pathway is crucial for the cellular response to bacteria [125] and viruses [126]. This pathway comprises three evolutionarily conserved proteins activated by phosphorylation: MAPKKKK, MAPKKK and MAPK [126]. In fish, three branches of the MAPK cascade have been identified: the extracellular signal-regulated kinase (ERK), the N-terminal c-Jun kinase (JNK) and the N-terminal c-Jun kinase (JNK). These have been shown to regulate various processes, including apoptosis, cell survival, differentiation and immune responses [127]. Although the upregulation of MAPK pathways would be anticipated given their role in the activation of inflammatory cytokines and active immune responses, a significant downregulation was observed in response to viruses and bacteria. This phenomenon has been documented in several studies in fish, including one on tilapia, in which the *map3k7* gene was found to be downregulated in response to tilapia lake virus. It is hypothesised that this mechanism may represent an evolutionary strategy employed by the virus to evade the immune response [128]. Moreover, *in vitro* studies indicate that the complete inhibition of MAPK/ERK function results in a reduction in viral load in TILV-infected cells, thereby suggesting that this pathway is indispensable for viral replication. These findings indicate that the downregulation of MAPK/ERK may facilitate the regulation of its RNA copies during the initial stages of infection [129]. Conversely, the activation or inhibition of MAPKKK pathways by bacteria has also been the subject of extensive study, as reviewed by Nandi and Aroeti (2023) [125]. Particularly in rainbow trout infected with *Vibrio anguillarum*, the *mkk1*, *mkk2a2* and *mkk4b3* genes were found to be upregulated, which may be indicative of a role in immunoregulation. However, *mkk6* was found to be downregulated, which could indicate negative feedback on MAPK activation due to overexpression of dual specificity phosphatases (DUSPs), which are key enzymes in MAPK negative feedback loops [130]. This study suggests a complex network where pathogens may co-evolve to survive after infection. Therefore, the *map2k6* and the rest of genes downregulated in this pathway (see TLR maps) emerge as a key element in our study that deserves further study for its potentially critical role in pathogen survival after infection.
