## Supplementary material for "Comparative transcriptomics of immune response to viral and bacterial stimuli in three acanthopterygian bony fish": Fig. S1

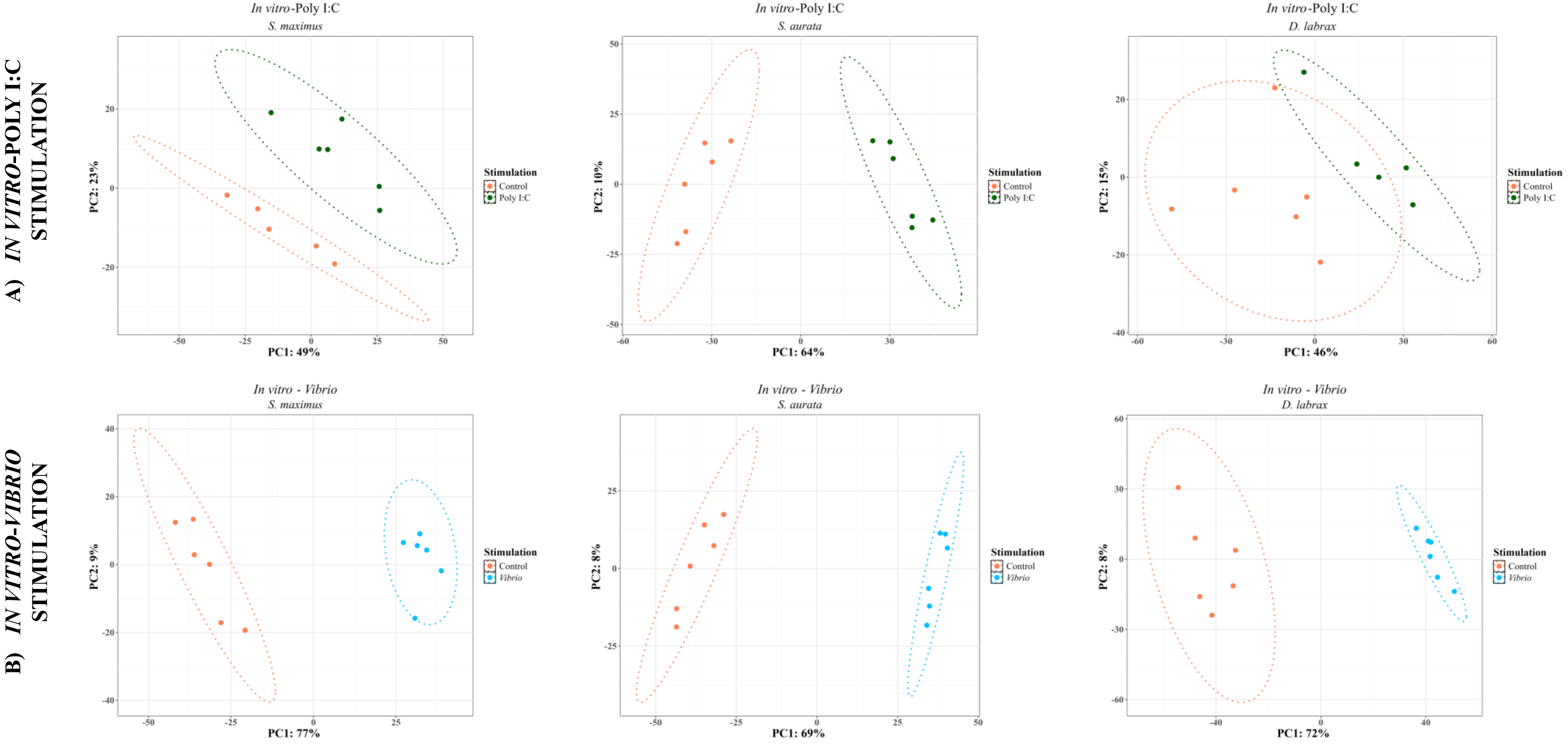
A) IN VITRO-POLY I:C STIMULATION
B) IN VITRO-VIBRIO STIMULATION
In vitro-Poly I:C
S. maximus
S. aurata
D. labrax
In vitro - Vibrio
PC1: 49%
PC2: 23%
PC1: 64%
PC2: 10%
PC1: 46%
PC2: 15%
PC1: 77%
PC2: 9%
PC1: 69%
PC2: 8%
PC1: 72%
PC2: 8%
Stimulation
Control
Poly I:C
Vibrio

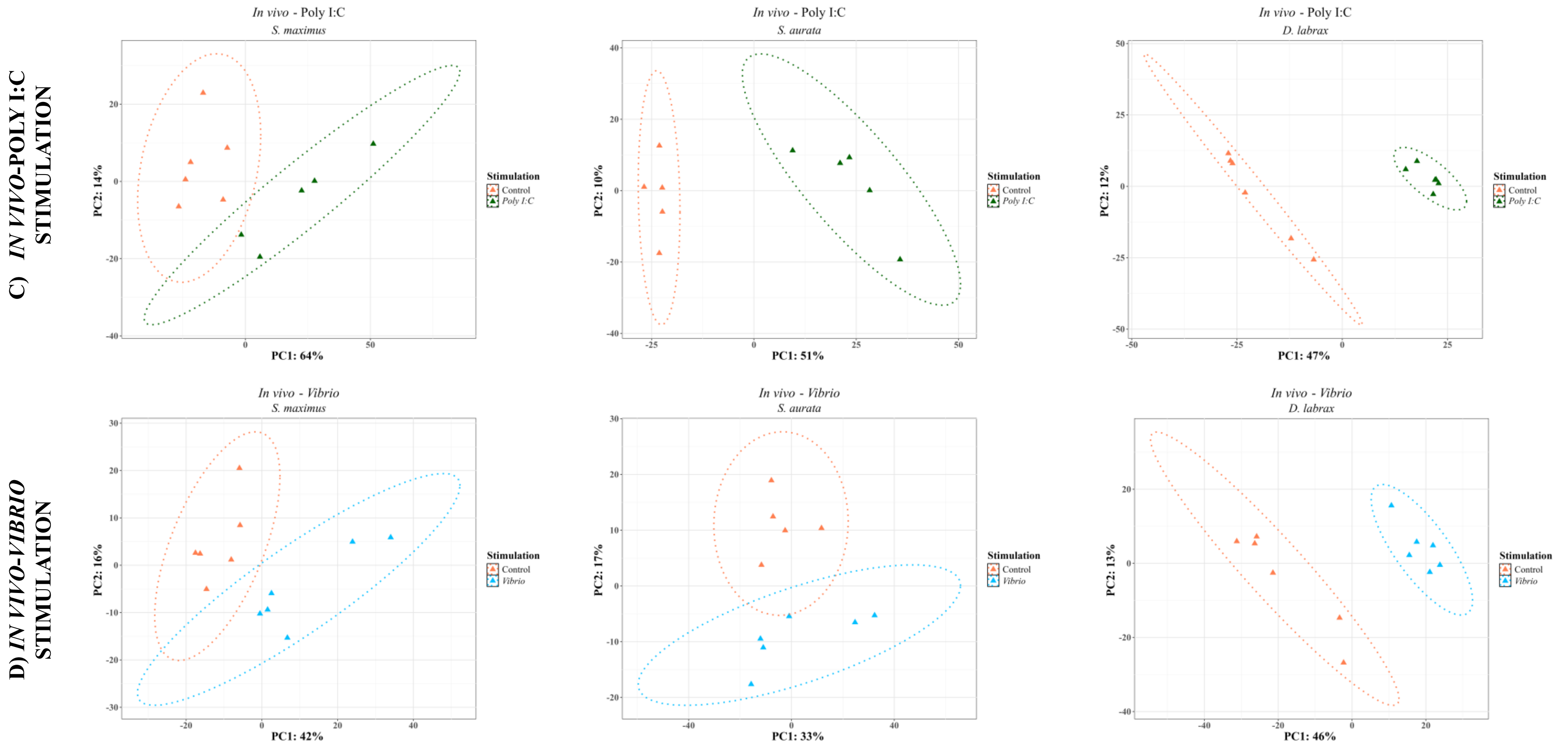


Fig S1. Principal Component Analysis of transcriptomic responses in *S. maximus, S. aurata* and *D. labrax* to Poly I:C (A, C) and *Vibrio* (B, D) stimulation *in vitro* and *in vivo* conditions. A) *In vitro*-Poly I:C stimulation revealed the greatest variance explained by PC1 in *S. aurata* (64%) and the lowest in *D. labrax* (46%), with PC2 (15%) enabling clear discrimination between the control and stimulated groups. B) *In Vitro-Vibrio* stimulation showed strong differentiation across species, with an average PC1 explained variance of 70%. C) In *in-vivo* conditions, Poly I:C stimulation resulted in PC1 explained variances ranging from 64% in *S. maximus* to 47% in *D. labrax*, with no overlap between control and stimulated samples across species. D) *In vivo-Vibrio* stimulation exhibited lower PC1 explained variance, particularly in *S. aurata* (33%), but higher PC2 explained variance (17%) allowed effective group discrimination, whereas *S. maximus* and *D. labrax* exhibited PC1 explained variances around 40% with sufficient explained variance by PC2 for discrimination.
