## Supplementary material for "Comparative transcriptomics of immune response to viral and bacterial stimuli in three acanthopterygian bony fish": Fig. S2

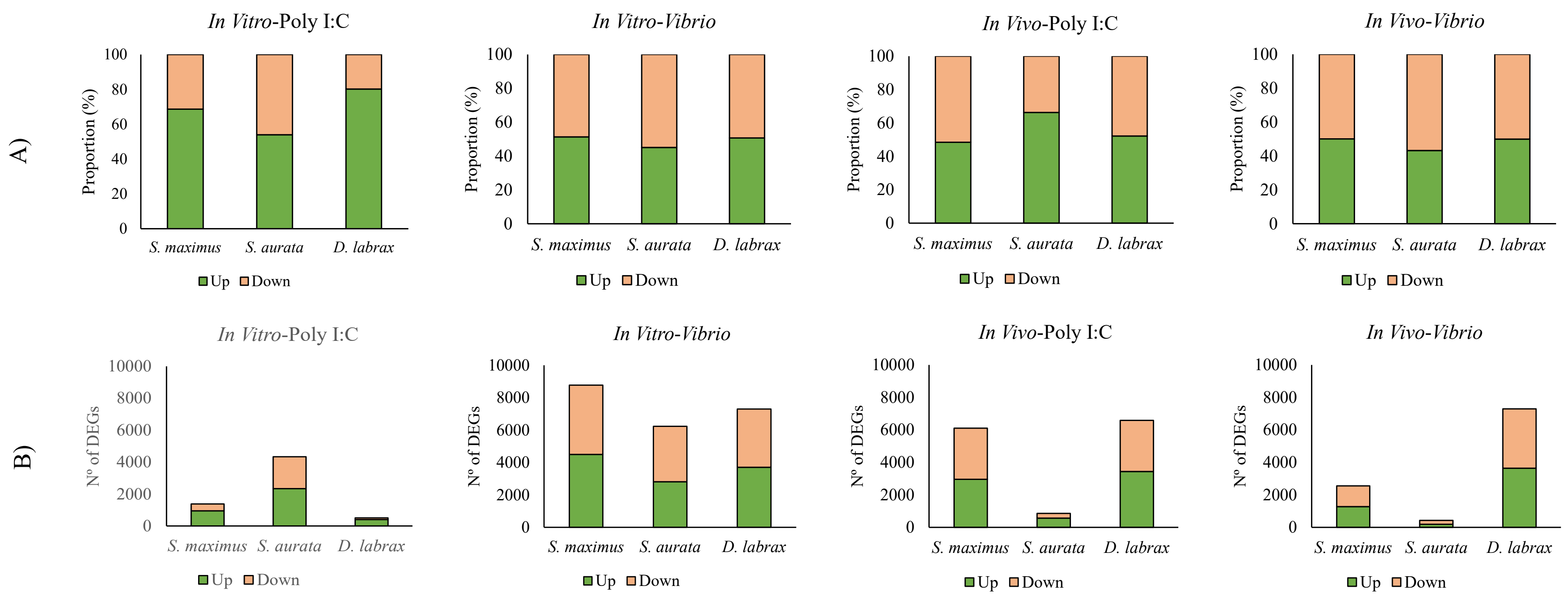


Fig S2. Proportion (A) and Number (B) of Up- and Down-regulated Differentially Expressed Genes (DEGs) in response to *Poly I:C* and *Vibrio* stimulation *In vitro* and *In vivo* in *S. maximus, S. aurata* and *D. labrax.*
