## Supplementary material for "Comparative transcriptomics of immune response to viral and bacterial stimuli in three acanthopterygian bony fish": Fig. S3

A)

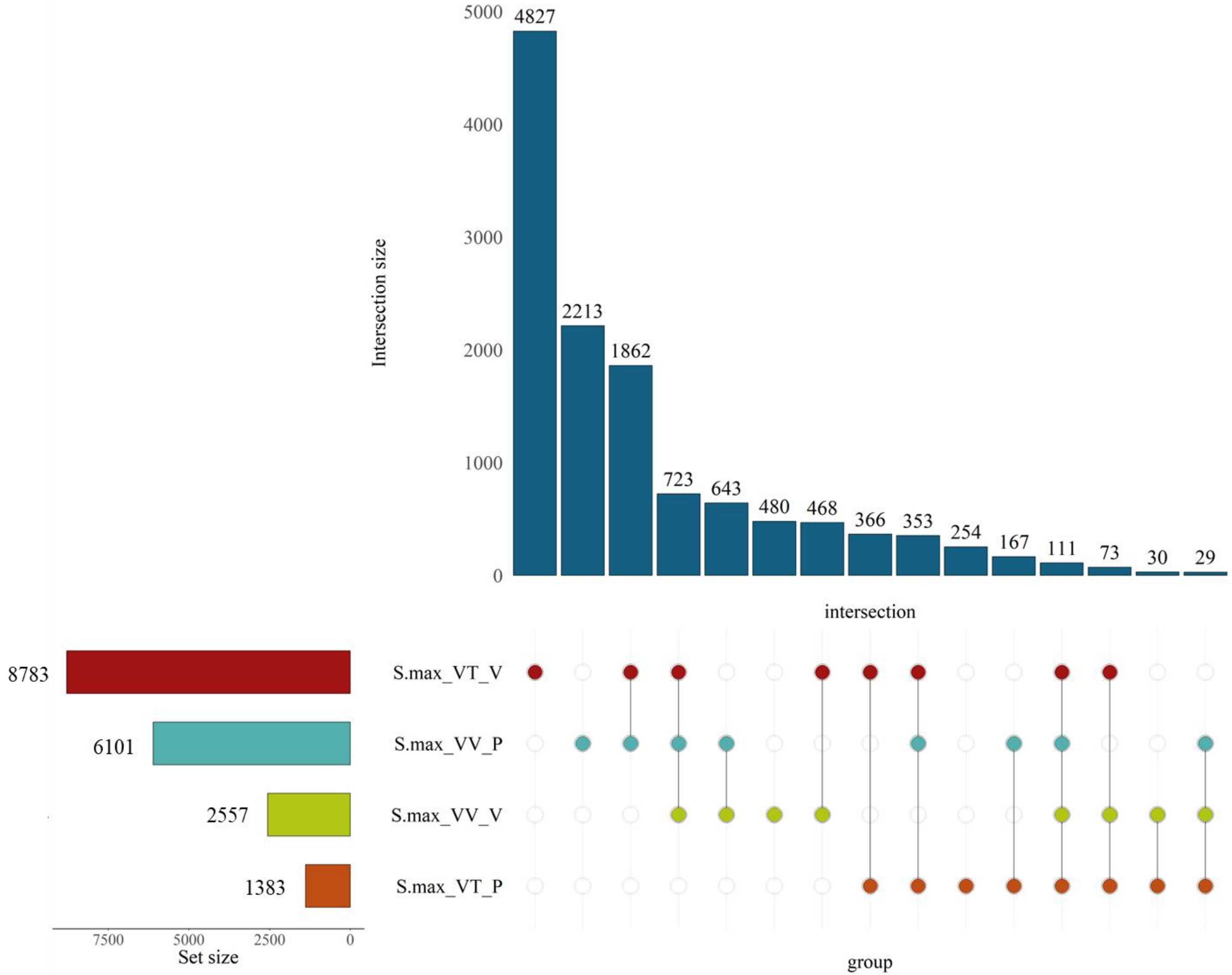

Intersection size
5000
4000
3000
2000
1000
0
4827
2213
1862
723
643
480
468
366
353
254
167
111
73
30
29
intersection
8783
6101
2557
1383
S.max_VT_V
S.max_VV_P
S.max_VV_V
S.max_VT_P
7500
5000
2500
0
Set size
group

B)

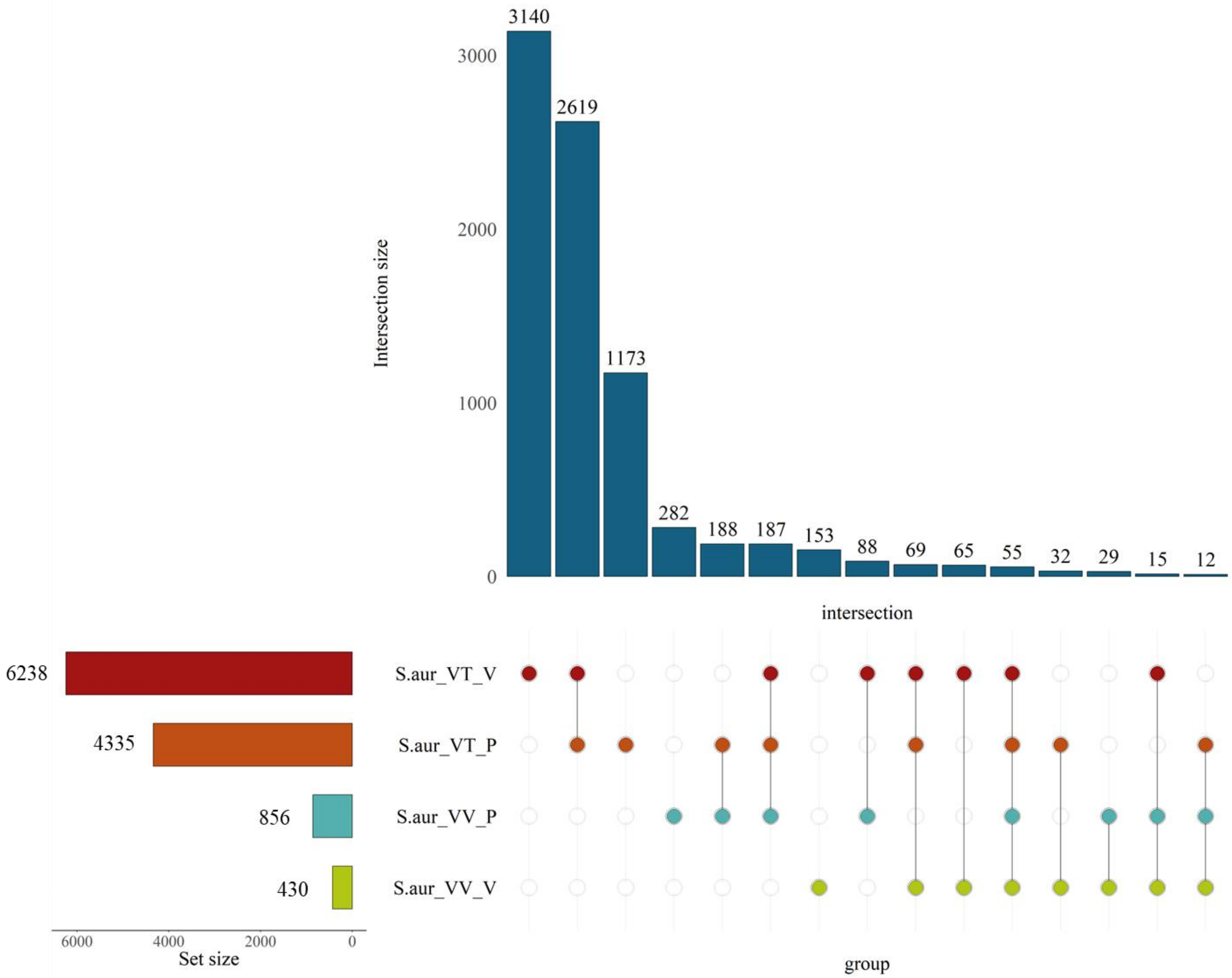

3140
2619
1173
282
188
187
153
88
69
65
55
32
29
15
12
3000
2000
1000
0
Intersection size
intersection
6238
4335
856
430
S.aur_VT_V
S.aur_VT_P
S.aur_VV_P
S.aur_VV_V
6000
4000
2000
0
Set size
group

C)

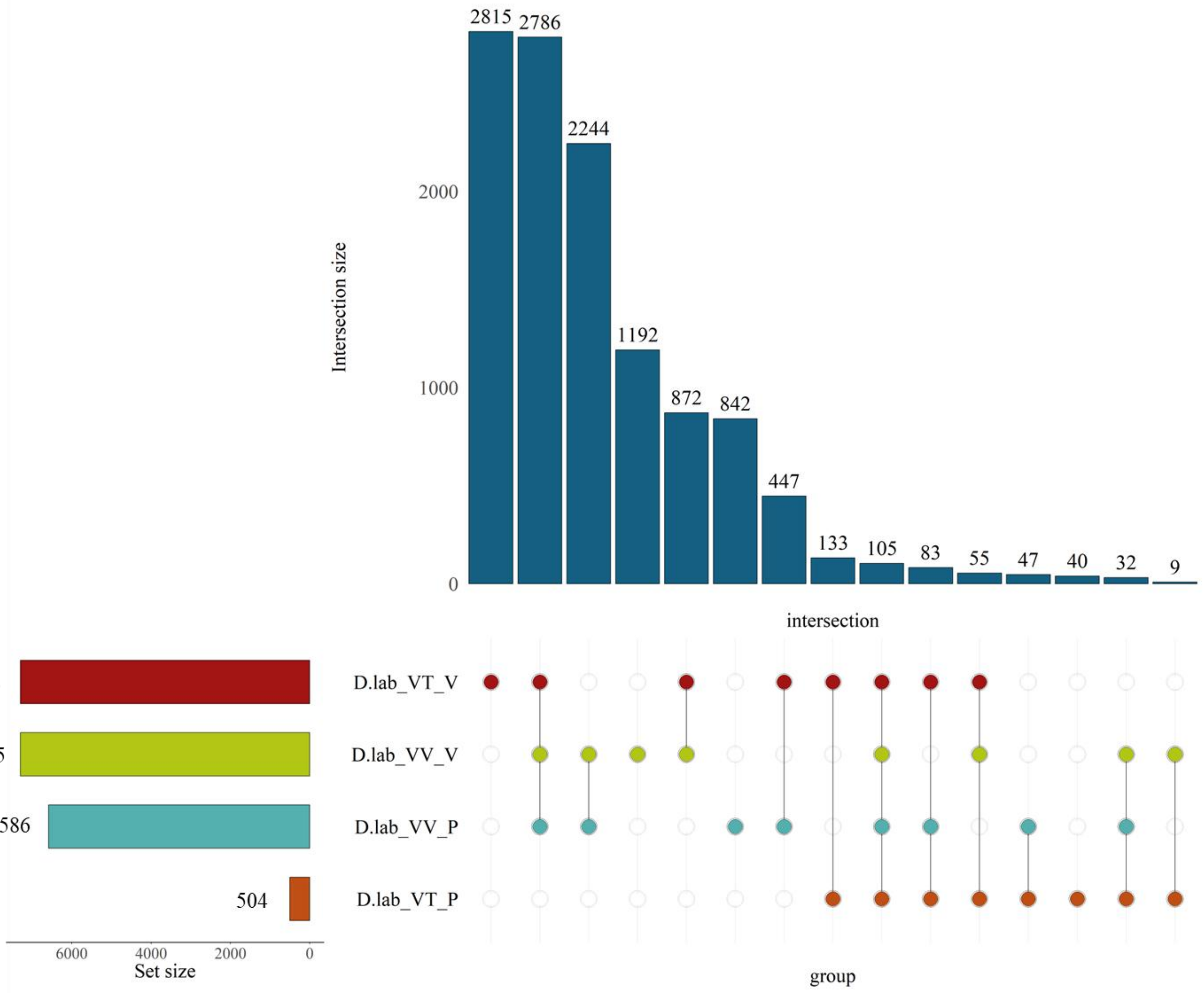

2815
2786
2244
1192
872
842
447
133
105
83
55
47
40
32
9
2000
1000
0
Intersection size
intersection
7296
7295
6586
504
D.lab_VT_V
D.lab_VV_V
D.lab_VV_P
D.lab_VT_P
6000
4000
2000
0
Set size
group

Fig S3. UpSet plots illustrating the overlap of DEGs for (A) *S. maximus*, (B) *S. aurata*, and (C) *D. labrax*. Vertical bar height represents the number of DEGs shared among specific combinations of experimental conditions. Horizontal bar widths represent the number of DEGs specific of each conditions: Red indicates *in vitro Vibrio* challenge (VT-V), orange *in vitro* Poly I:C stimulation (VT-P), blue *in vivo* Poly I:C challenge (VV-P), and green *in vivo Vibrio* challenge (VV-V). Coloured set size bars and intersection dots correspond to each condition. Numbers above each bar indicate the intersection size of DEGs.
