## Supplementary material for "Comparative transcriptomics of immune response to viral and bacterial stimuli in three acanthopterygian bony fish": Fig. S4

A)

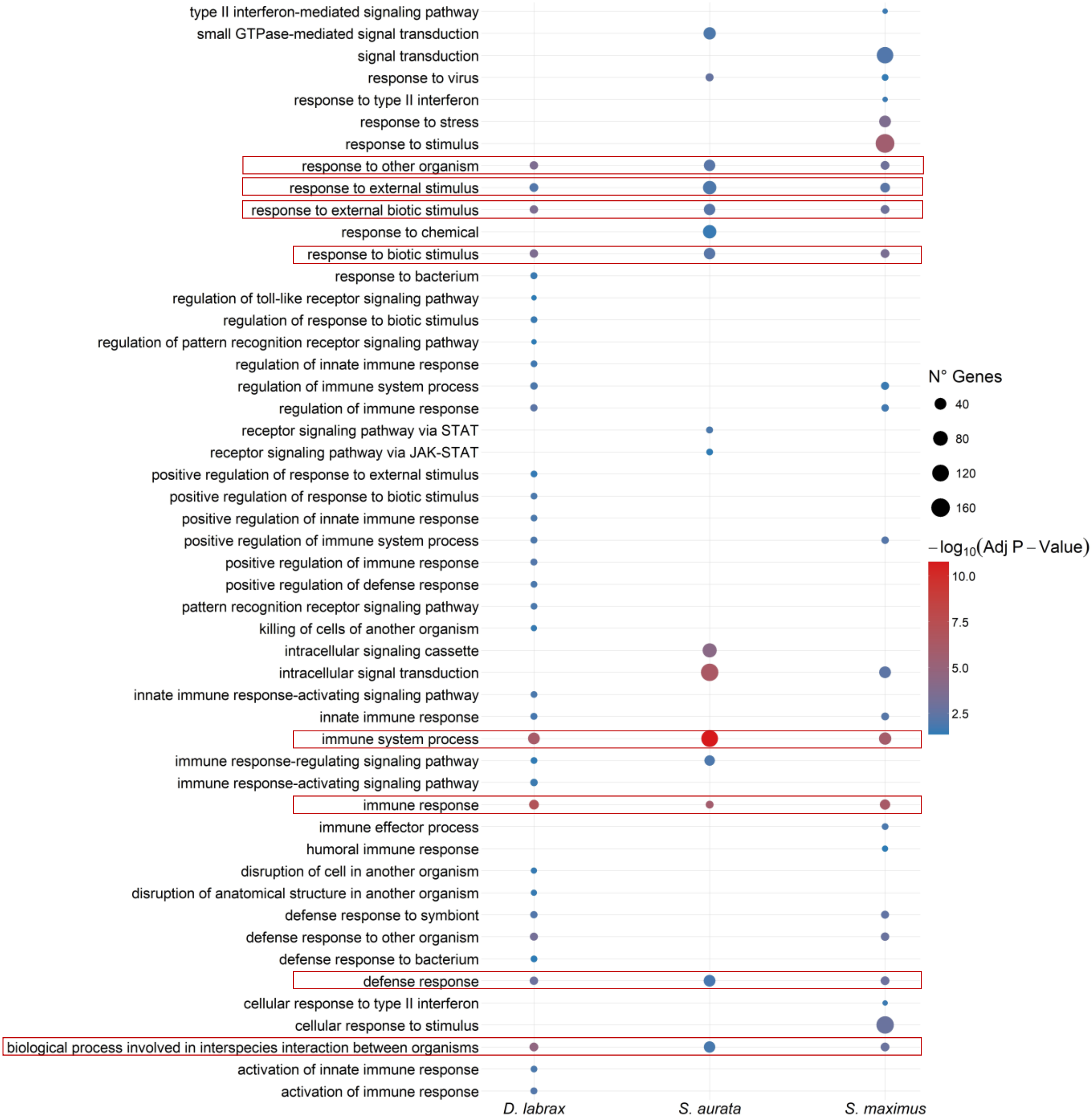

type II interferon-mediated signaling pathway
small GTPase-mediated signal transduction
signal transduction
response to virus
response to type II interferon
response to stress
response to stimulus
response to other organism
response to external stimulus
response to external biotic stimulus
response to chemical
response to biotic stimulus
response to bacterium
regulation of toll-like receptor signaling pathway
regulation of response to biotic stimulus
regulation of pattern recognition receptor signaling pathway
regulation of innate immune response
regulation of immune system process
regulation of immune response
receptor signaling pathway via STAT
receptor signaling pathway via JAK-STAT
positive regulation of response to external stimulus
positive regulation of response to biotic stimulus
positive regulation of innate immune response
positive regulation of immune system process
positive regulation of immune response
positive regulation of defense response
pattern recognition receptor signaling pathway
killing of cells of another organism
intracellular signaling cassette
intracellular signal transduction
innate immune response-activating signaling pathway
innate immune response
immune system process
immune response-regulating signaling pathway
immune response-activating signaling pathway
immune response
immune effector process
humoral immune response
disruption of cell in another organism
disruption of anatomical structure in another organism
defense response to symbiont
defense response to other organism
defense response to bacterium
defense response
cellular response to type II interferon
cellular response to stimulus
biological process involved in interspecies interaction between organisms
activation of innate immune response
activation of immune response
D. labrax
S. aurata
S. maximus
N° Genes
40
80
120
160
$-\log_{10}$(Adj P – Value)
10.0
7.5
5.0
2.5

B)

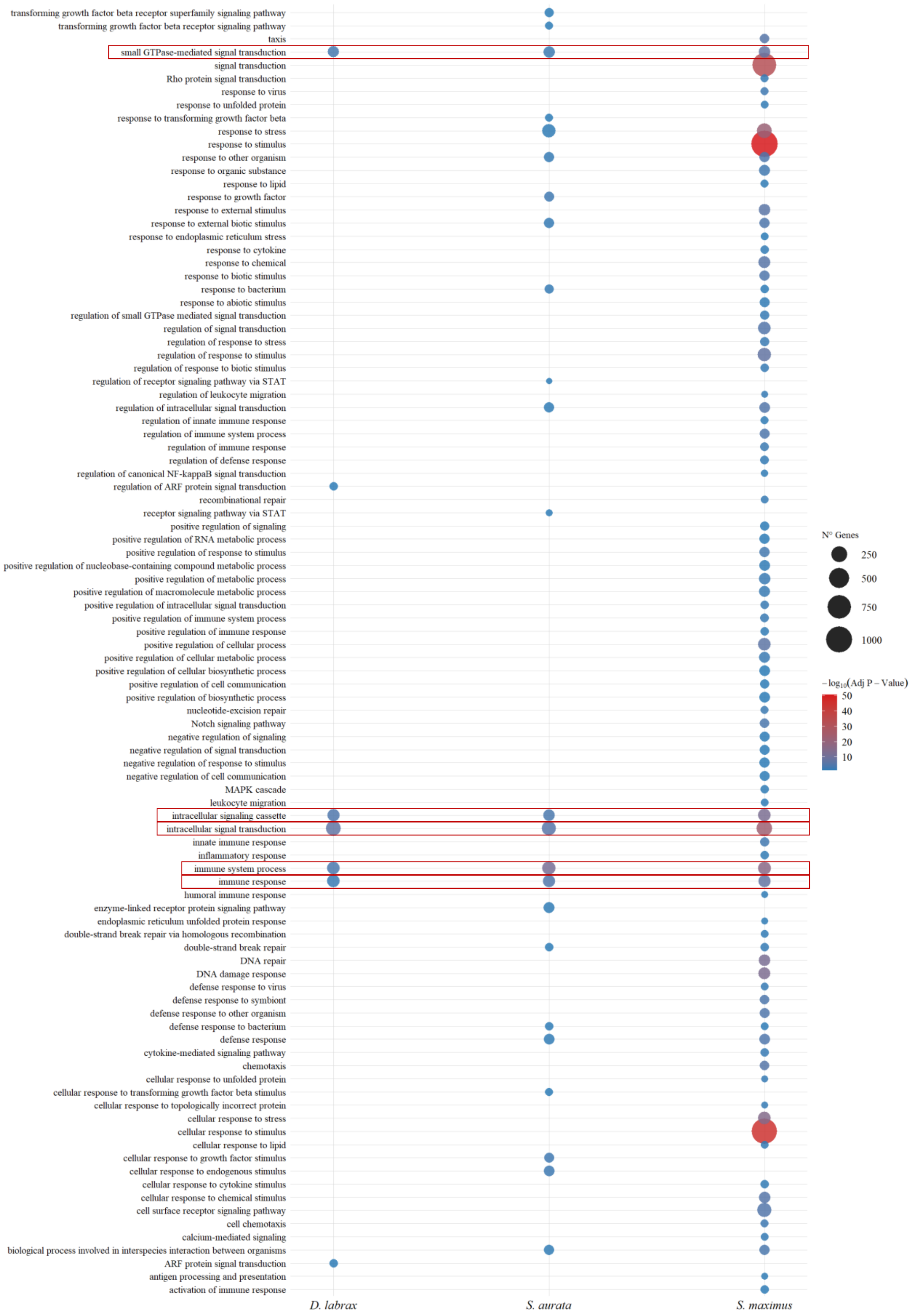

transforming growth factor beta receptor superfamily signaling pathway
transforming growth factor beta receptor signaling pathway
taxis
small GTPase-mediated signal transduction
signal transduction
Rho protein signal transduction
response to virus
response to unfolded protein
response to transforming growth factor beta
response to stress
response to stimulus
response to other organism
response to organic substance
response to lipid
response to growth factor
response to external stimulus
response to external biotic stimulus
response to endoplasmic reticulum stress
response to cytokine
response to chemical
response to biotic stimulus
response to bacterium
response to abiotic stimulus
regulation of small GTPase mediated signal transduction
regulation of signal transduction
regulation of response to stress
regulation of response to stimulus
regulation of response to biotic stimulus
regulation of receptor signaling pathway via STAT
regulation of leukocyte migration
regulation of intracellular signal transduction
regulation of innate immune response
regulation of immune system process
regulation of immune response
regulation of defense response
regulation of canonical NF-kappaB signal transduction
regulation of ARF protein signal transduction
recombinational repair
receptor signaling pathway via STAT
positive regulation of signaling
positive regulation of RNA metabolic process
positive regulation of response to stimulus
positive regulation of nucleobase-containing compound metabolic process
positive regulation of metabolic process
positive regulation of macromolecule metabolic process
positive regulation of intracellular signal transduction
positive regulation of immune system process
positive regulation of immune response
positive regulation of cellular process
positive regulation of cellular metabolic process
positive regulation of cellular biosynthetic process
positive regulation of cell communication
positive regulation of biosynthetic process
nucleotide-excision repair
Notch signaling pathway
negative regulation of signaling
negative regulation of signal transduction
negative regulation of response to stimulus
negative regulation of cell communication
MAPK cascade
leukocyte migration
intracellular signaling cassette
intracellular signal transduction
innate immune response
inflammatory response
immune system process
immune response
humoral immune response
enzyme-linked receptor protein signaling pathway
endoplasmic reticulum unfolded protein response
double-strand break repair via homologous recombination
double-strand break repair
DNA repair
DNA damage response
defense response to virus
defense response to symbiont
defense response to other organism
defense response to bacterium
defense response
cytokine-mediated signaling pathway
chemotaxis
cellular response to unfolded protein
cellular response to transforming growth factor beta stimulus
cellular response to topologically incorrect protein
cellular response to stress
cellular response to stimulus
cellular response to lipid
cellular response to growth factor stimulus
cellular response to endogenous stimulus
cellular response to cytokine stimulus
cellular response to chemical stimulus
cell surface receptor signaling pathway
cell chemotaxis
calcium-mediated signaling
biological process involved in interspecies interaction between organisms
ARF protein signal transduction
antigen processing and presentation
activation of immune response
D. labrax
S. aurata
S. maximus
N° Genes
250
500
750
1000
−log10(Adj P − Value)
50
40
30
20
10

C)

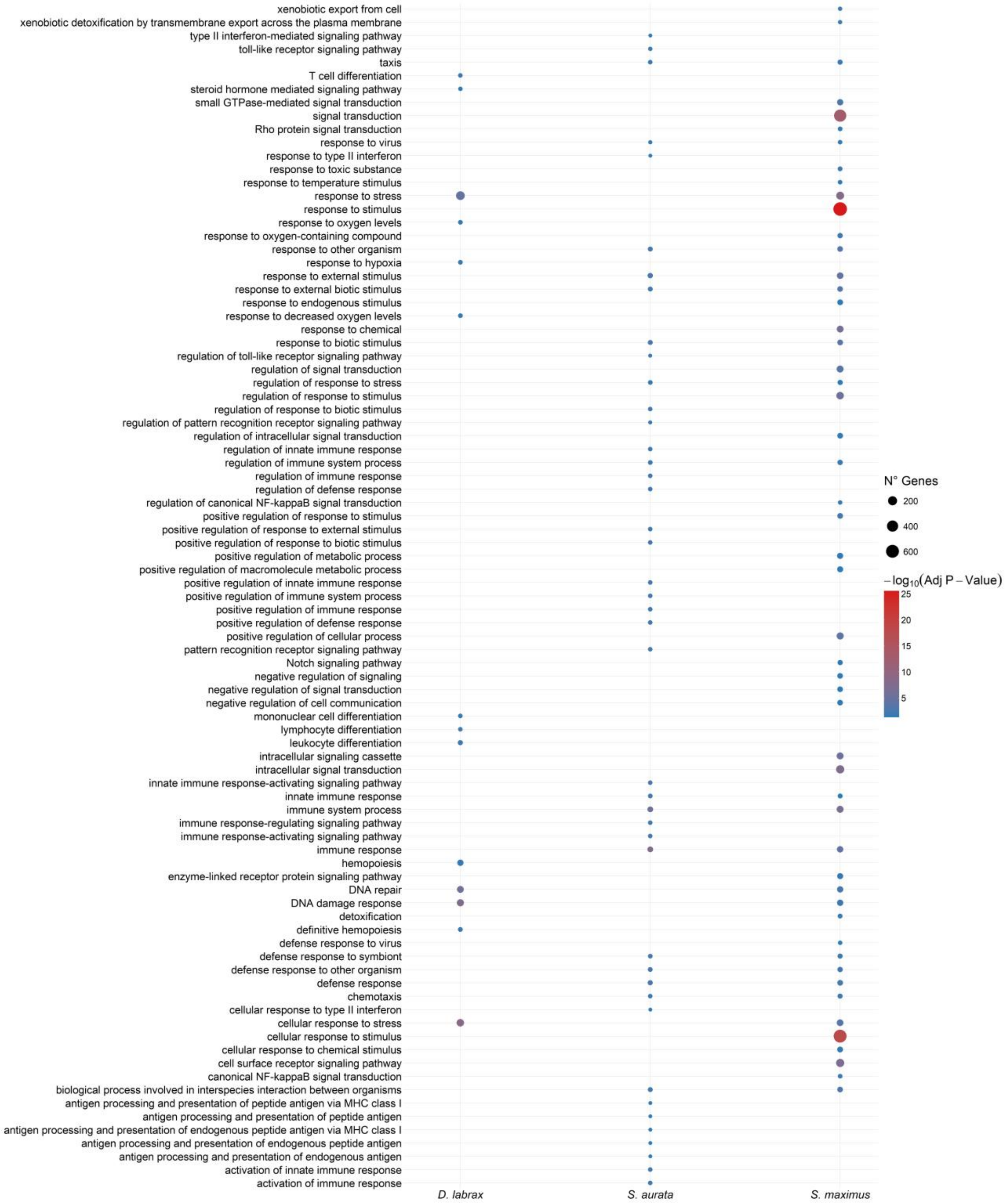
xenobiotic export from cell
xenobiotic detoxification by transmembrane export across the plasma membrane
type II interferon-mediated signaling pathway
toll-like receptor signaling pathway
taxis
T cell differentiation
steroid hormone mediated signaling pathway
small GTPase-mediated signal transduction
signal transduction
Rho protein signal transduction
response to virus
response to type II interferon
response to toxic substance
response to temperature stimulus
response to stress
response to stimulus
response to oxygen levels
response to oxygen-containing compound
response to other organism
response to hypoxia
response to external stimulus
response to external biotic stimulus
response to endogenous stimulus
response to decreased oxygen levels
response to chemical
response to biotic stimulus
regulation of toll-like receptor signaling pathway
regulation of signal transduction
regulation of response to stress
regulation of response to stimulus
regulation of response to biotic stimulus
regulation of pattern recognition receptor signaling pathway
regulation of intracellular signal transduction
regulation of innate immune response
regulation of immune system process
regulation of immune response
regulation of defense response
regulation of canonical NF-kappaB signal transduction
positive regulation of response to stimulus
positive regulation of response to external stimulus
positive regulation of response to biotic stimulus
positive regulation of metabolic process
positive regulation of macromolecule metabolic process
positive regulation of innate immune response
positive regulation of immune system process
positive regulation of immune response
positive regulation of defense response
positive regulation of cellular process
pattern recognition receptor signaling pathway
Notch signaling pathway
negative regulation of signaling
negative regulation of signal transduction
negative regulation of cell communication
mononuclear cell differentiation
lymphocyte differentiation
leukocyte differentiation
intracellular signaling cassette
intracellular signal transduction
innate immune response-activating signaling pathway
innate immune response
immune system process
immune response-regulating signaling pathway
immune response-activating signaling pathway
immune response
hemopoiesis
enzyme-linked receptor protein signaling pathway
DNA repair
DNA damage response
detoxification
definitive hemopoiesis
defense response to virus
defense response to symbiont
defense response to other organism
defense response
chemotaxis
cellular response to type II interferon
cellular response to stress
cellular response to stimulus
cellular response to chemical stimulus
cell surface receptor signaling pathway
canonical NF-kappaB signal transduction
biological process involved in interspecies interaction between organisms
antigen processing and presentation of peptide antigen via MHC class I
antigen processing and presentation of peptide antigen
antigen processing and presentation of endogenous peptide antigen via MHC class I
antigen processing and presentation of endogenous peptide antigen
antigen processing and presentation of endogenous antigen
activation of innate immune response
activation of immune response
D. labrax
S. aurata
S. maximus
N° Genes
200
400
600
$-\log_{10}$(Adj P − Value)
25
20
15
10
5

D)

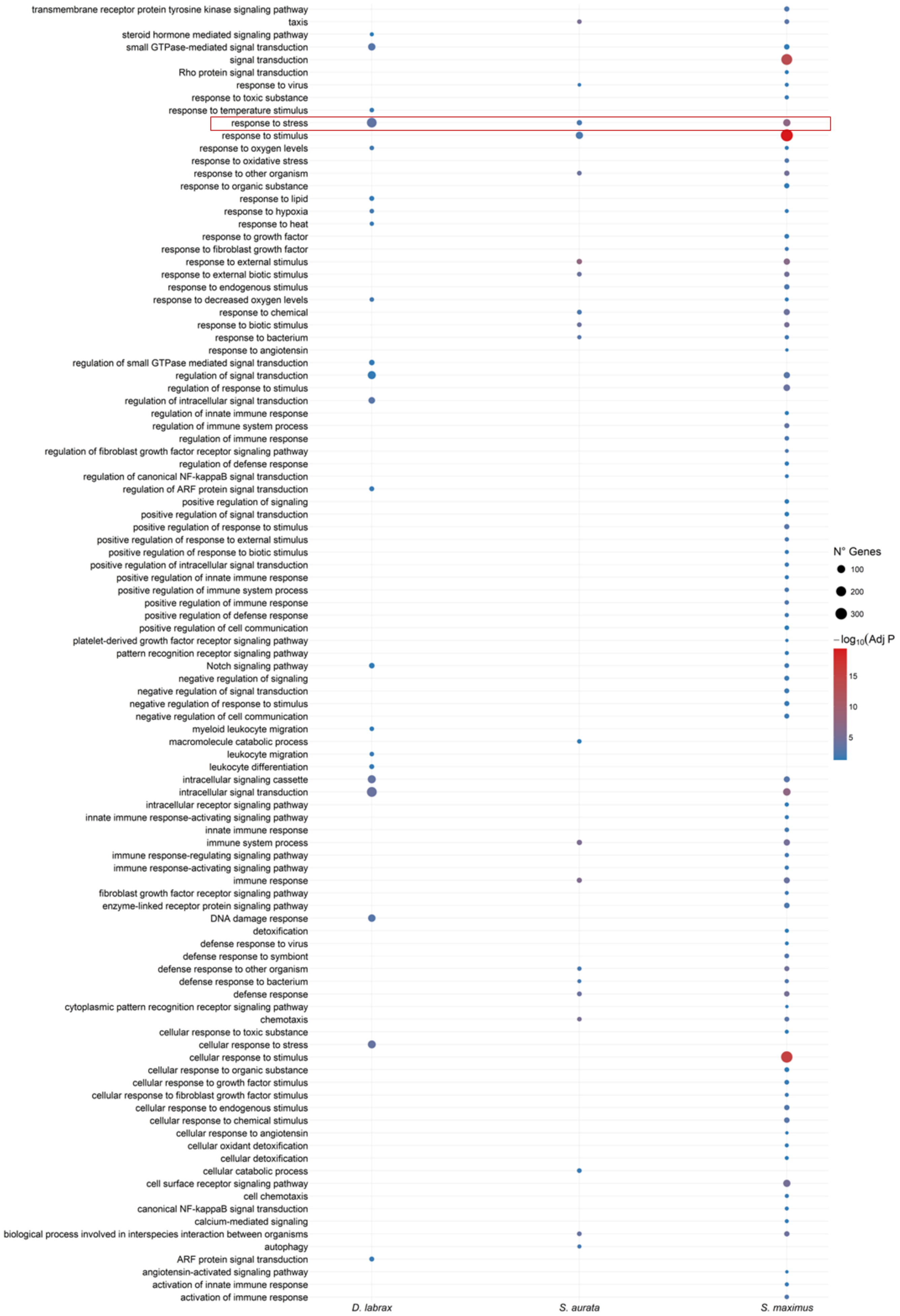
transmembrane receptor protein tyrosine kinase signaling pathway
taxis
steroid hormone mediated signaling pathway
small GTPase-mediated signal transduction
signal transduction
Rho protein signal transduction
response to virus
response to toxic substance
response to temperature stimulus
response to stress
response to stimulus
response to oxygen levels
response to oxidative stress
response to other organism
response to organic substance
response to lipid
response to hypoxia
response to heat
response to growth factor
response to fibroblast growth factor
response to external stimulus
response to external biotic stimulus
response to endogenous stimulus
response to decreased oxygen levels
response to chemical
response to biotic stimulus
response to bacterium
response to angiotensin
regulation of small GTPase mediated signal transduction
regulation of signal transduction
regulation of response to stimulus
regulation of intracellular signal transduction
regulation of innate immune response
regulation of immune system process
regulation of immune response
regulation of fibroblast growth factor receptor signaling pathway
regulation of defense response
regulation of canonical NF-kappaB signal transduction
regulation of ARF protein signal transduction
positive regulation of signaling
positive regulation of signal transduction
positive regulation of response to stimulus
positive regulation of response to external stimulus
positive regulation of response to biotic stimulus
positive regulation of intracellular signal transduction
positive regulation of innate immune response
positive regulation of immune system process
positive regulation of immune response
positive regulation of defense response
positive regulation of cell communication
platelet-derived growth factor receptor signaling pathway
pattern recognition receptor signaling pathway
Notch signaling pathway
negative regulation of signaling
negative regulation of signal transduction
negative regulation of response to stimulus
negative regulation of cell communication
myeloid leukocyte migration
macromolecule catabolic process
leukocyte migration
leukocyte differentiation
intracellular signaling cassette
intracellular signal transduction
intracellular receptor signaling pathway
innate immune response-activating signaling pathway
innate immune response
immune system process
immune response-regulating signaling pathway
immune response-activating signaling pathway
immune response
fibroblast growth factor receptor signaling pathway
enzyme-linked receptor protein signaling pathway
DNA damage response
detoxification
defense response to virus
defense response to symbiont
defense response to other organism
defense response to bacterium
defense response
cytoplasmic pattern recognition receptor signaling pathway
chemotaxis
cellular response to toxic substance
cellular response to stress
cellular response to stimulus
cellular response to organic substance
cellular response to growth factor stimulus
cellular response to fibroblast growth factor stimulus
cellular response to endogenous stimulus
cellular response to chemical stimulus
cellular response to angiotensin
cellular oxidant detoxification
cellular detoxification
cellular catabolic process
cell surface receptor signaling pathway
cell chemotaxis
canonical NF-kappaB signal transduction
calcium-mediated signaling
biological process involved in interspecies interaction between organisms
autophagy
ARF protein signal transduction
angiotensin-activated signaling pathway
activation of innate immune response
activation of immune response
D. labrax
S. aurata
S. maximus
N° Genes
100
200
300
−log10(Adj P
15
10
5

Fig. S4. Bubble plots display significantly enriched GO terms filtered through REVIGO and associated with DEGs for *S. maximus* (right), *S. aurata* (centre), and *D. labrax* (left) under four experimental conditions: A) *In vitro* response to Poly I:C stimulation; B) *In vitro* response to *Vibrio* stimulation; C) *In vivo* response to Poly I:C stimulation; D) *In vivo* response to *Vibrio* stimulation. The size of each bubble represents the number of genes associated with the corresponding GO term, where larger bubbles indicate greater gene involvement in the biological process. The colour gradient represents the statistical significance expressed as $-\log_{10}$ of the adjusted P-value, from blue (lower significance) to red (higher significance). GO terms enclosed in red squares represent biological processes shared among the three species.
