## Supplementary material for "Comparative transcriptomics of immune response to viral and bacterial stimuli in three acanthopterygian bony fish": Fig. S5

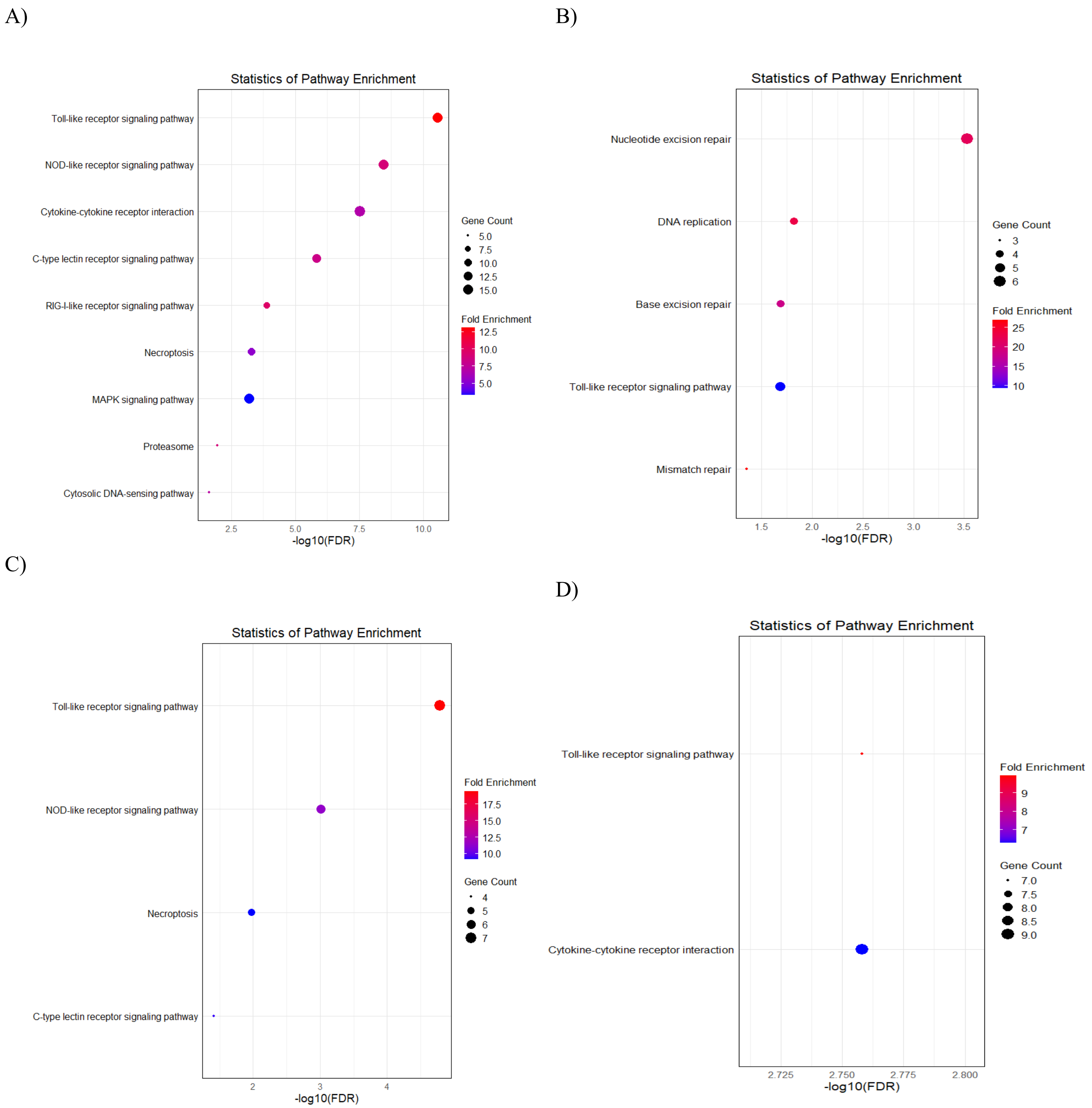


Fig. S5. KEGG related with immune system *in vitro* (A) and *in vivo* (B), with Poly I:C (C) and with *Vibrio* (D). The analysis used differentially expressed genes in at least one of the three species and maintaining consistent regulation direction in their orthologs. Data obtained from Table S10B (*in vitro*), Table S10C (*in vivo*), Table S10D (Poly I:C) and Table S10E (*Vibrio*).
