## Supplementary material for "Comparative transcriptomics of immune response to viral and bacterial stimuli in three acanthopterygian bony fish": Table S8

**Table S8**. Differentially expressed orthologous genes between control and Poly I:C stimulation in *S. maximus, S. aurata* and *D. labrax*: Comparison of *In Vitro -In Vivo* responses.

| **Symbol ID*** | ***S. maximus* Gen ID** | ***S. maximus*** | | ***S. aurata* Gen ID** | ***S. aurata*** | | ***D. labrax* Gen ID** | ***D. labrax*** | | |
|---|---|---|---|---|---|---|---|---|---|---|
| | | ***In Vitro* Fold Change** | ***In Vivo* Fold Change** | | ***In Vitro* Fold Change** | ***In Vivo* Fold Change** | | ***In Vitro* Fold Change** | ***In Vivo* Fold Change** | |
| BPIFCL | *ENSSMAG00000015055* | 2.35 | 0.59 | *ENSSAUG00010011515* | 1.33 | 1.05 | *ENSDLAG00005016879* | 1.13 | 0.80 | 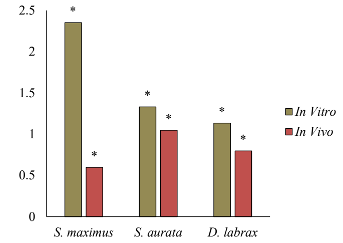 |
| IFI35 | *ENSSMAG00000007822* | 1.33 | 1.10 | *ENSSAUG00010018946* | 1.20 | 1.62 | *ENSDLAG00005021869* | 0.93 | 01.23 | 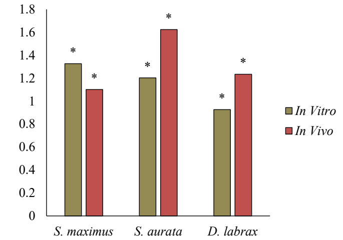 |
| IFI44 | *ENSSMAG00000021010* | 1.72 | 0.91 | *ENSSAUG00010009655* | 4.17 | 4.10 | *ENSDLAG00005019782* | 1.67 | 3.38 | 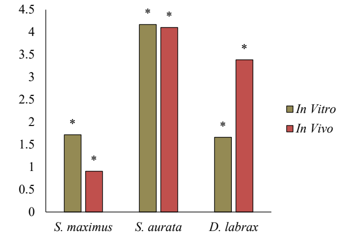 |
| IFIH1 | *ENSSMAG00000012345* | 1.23 | 0.62 | *ENSSAUG00010021326* | 3.60 | 4.27 | *ENSDLAG00005019389* | 1.14 | 2.01 | 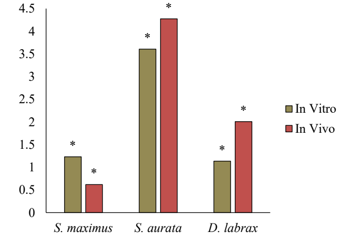 |

| | | | | | | | | | | |
|---|---|---|---|---|---|---|---|---|---|---|
| LGALS9L | *ENSSMAG00000015775* | 1.05 | 0.80 | *ENSSAUG00010012079* | 1.73 | 1.20 | *ENSDLAG00005001138* | 0.63 | 1.32 | 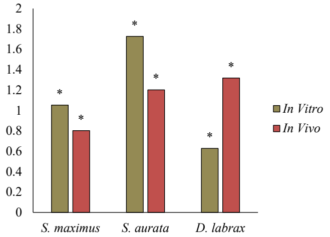 |
| NLRC5 | *ENSSMAG00000014651* | 1.85 | 1.38 | *ENSSAUG00010012287* | 1.75 | 2.57 | *ENSDLAG00005032672* | 1.47 | 1.66 | 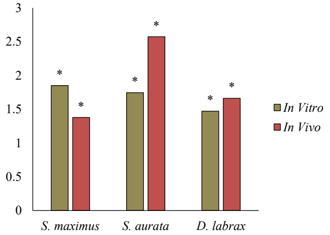 |
| STAT1a | ENSSMAG00000012718 | 1.56 | 0.80 | *ENSSAUG00010021056* | 2.32 | 2.50 | *ENSDLAG00005003574* | 2.40 | 2.32 | 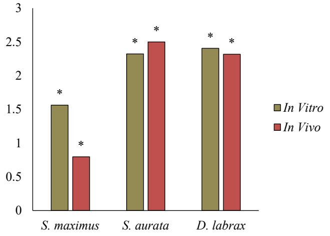 |

* The Symbol ID represents the most common Ensembl gene name found across the species studied.

Asterisks in the graph (*) indicate significant differences (p<0.05) between control and stimulated conditions for each species in individual experiments (Data: Table S7).
