## Supplementary material for "Comparative transcriptomics of immune response to viral and bacterial stimuli in three acanthopterygian bony fish": Table S9

**Table S9**. Differentially expressed orthologous genes between control-Poly I:C and control-*Vibrio* stimulation in *S. maximus, S. aurata* and *D. labrax*: Comparison of *In Vitro* model.

| **Symbol ID*** | ***S. maximus* Gen ID** | ***S. maximus*** | | ***S. aurata* Gen ID** | ***S. aurata*** | | ***D. labrax* Gen ID** | ***D. labrax*** | | |
|---|---|---|---|---|---|---|---|---|---|---|
| | | **Poly I:C Fold Change** | **Vibrio Fold Change** | | **Poly I:C Fold Change** | ***Vibrio* Fold Change** | | **Poly I:C Fold Change** | ***Vibrio* Fold Change** | |
| BPIFCL | *ENSSMAG00000015055* | 2.35 | -0.86 | *ENSSAUG00010011515* | 1.33 | -1.53 | *ENSDLAG00005016879* | 1.13 | -1.80 | 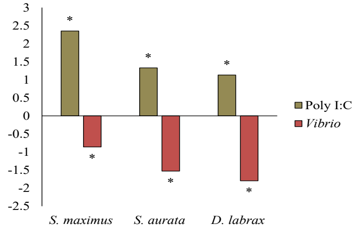  |
| CCL19 | *ENSSMAG00000034610* | 4.32 | 2.20 | *ENSSAUG00010021807* | 5.81 | 4.90 | *ENSDLAG00005033541* | 2.62 | 2.40 | 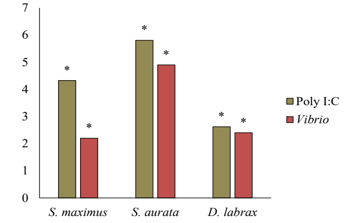  |
| IFI35 | *ENSSMAG00000007822* | 1.33 | 1.25 | *ENSSAUG00010018946* | 1.20 | 1.04 | *ENSDLAG00005021869* | 0.93 | 0.93 | 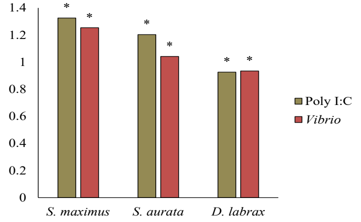  |
| IRF3 | *ENSSMAG00000015932* | 1.81 | 2.20 | *ENSSAUG00010024935* | 3.67 | 1.78 | *ENSDLAG00005013577* | 3.63 | 2.49 | 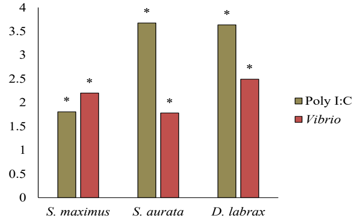  |

| | | | | | | | | | | |
|---|---|---|---|---|---|---|---|---|---|---|
| LY75 | *ENSSMAG00000012324* | 1.28 | 1.59 | ENSSAUG00010022226 | 0.99 | -1.17 | *ENSDLAG00005014934* | 0.59 | -0.68 | 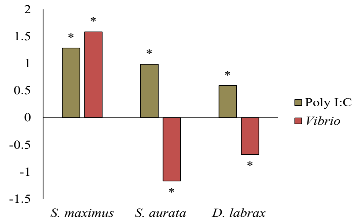 |
| ISG15 | *ENSSMAG00000014327* | 3.43 | 0.85 | *ENSSAUG00010020050* | 5.39 | 2.08 | *ENSDLAG00005035060* | 6.72 | 5.78 | 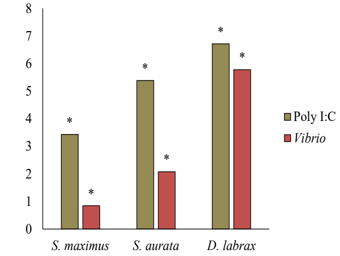 |
| PSME1 | *ENSSMAG00000036525* | 0.48 | 1.23 | *ENSSAUG00010011610* | 1.54 | 2.02 | *ENSDLAG00005004361* | 0.95 | 1.72 | 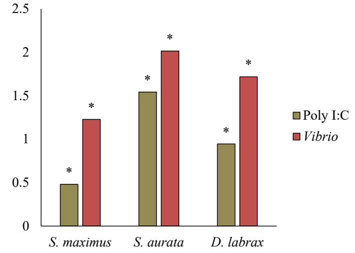 |
| PSME2 | *ENSSMAG00000007870* | 0.67 | 1.25 | *ENSSAUG00010024593* | 1.62 | 2.49 | *ENSDLAG00005030547* | 1.20 | 1.55 | 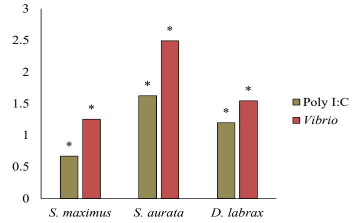 |

| | | | | | | | | | | |
|---|---|---|---|---|---|---|---|---|---|---|
| PSMB13a | *ENSSMAG00000017232* | 0.35 | 0.60 | *ENSSAUG00010020621* | 0.96 | 1.47 | *ENSDLAG00005033124* | 0.82 | 0.84 | 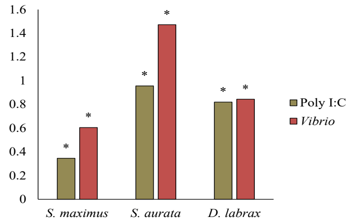  |
| SOCS1b | *ENSSMAG00000002682* | 1.98 | 2.40 | *ENSSAUG00010009323* | 2.06 | 1.81 | *ENSDLAG00005033441* | 1.26 | 0.79 | 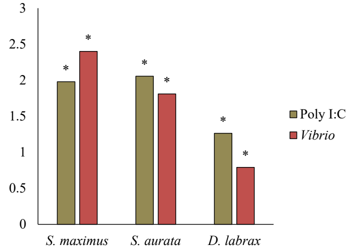  |
| TAPBP.2 | *ENSSMAG00000017420* | 1.04 | 1.12 | *ENSSAUG00010000163* | 1.44 | 1.27 | *ENSDLAG00005003462* | 1.01 | 0.85 | 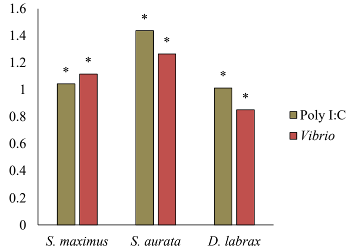  |
| ETV7 | *ENSSMAG00000006052* | 1.03 | 0.61 | *ENSSAUG00010004327* | 1.95 | 1.38 | *ENSDLAG00005019610* | 1.27 | 1.05 | 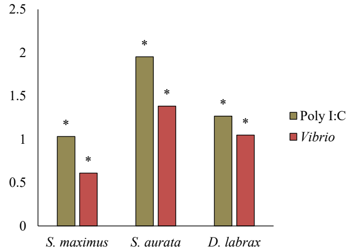  |

| | | | | | | | | | | |
|---|---|---|---|---|---|---|---|---|---|---|
| TAP1 | *ENSSMAG00000013604* | 0.99 | 2.14 | *ENSSAUG00010015502* | 1.96 | 2.82 | *ENSDLAG00005006626* | 0.98 | 0.71 | 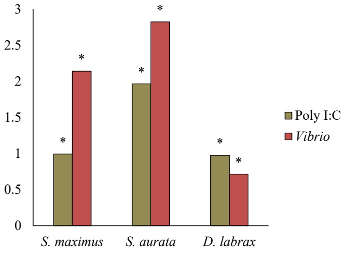 |
| TAP2a | *ENSSMAG00000016419* | 1.35 | 0.51 | *ENSSAUG00010020600* | 3.29 | 3.16 | *ENSDLAG00005012408* | 1.36 | 1.44 | 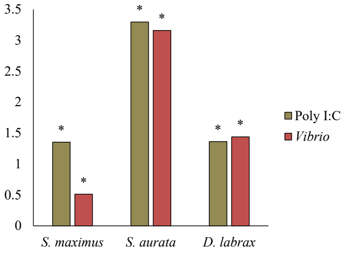 |
| UBA7 | *ENSSMAG00000016945* | 1.27 | 0.53 | *ENSSAUG00010020707* | 3.88 | 2.89 | *ENSDLAG00005023600* | 2.45 | 1.52 | 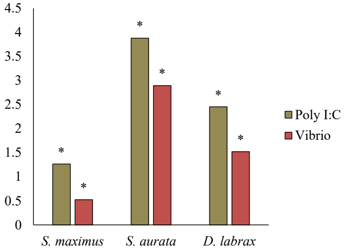 |

* The Symbol ID represents the most common Ensembl gene name found across the species studied.

Asterisks in the graph (*) indicate significant differences ($p<0.05$) between control and stimulated conditions for each species in individual experiments (Data: Table S7).
